# Native RNA nanopore sequencing reveals antibiotic-induced loss of rRNA modifications in the A- and P-sites

**DOI:** 10.1101/2023.03.21.533606

**Authors:** Anna Delgado-Tejedor, Rebeca Medina, Oguzhan Begik, Luca Cozzuto, Julia Ponomarenko, Eva Maria Novoa

**Affiliations:** Centre for Genomic Regulation (CRG), The Barcelona Institute of Science and Technology, Dr. Aiguader 88, Barcelona 08003, Spain; Universitat Pompeu Fabra, Barcelona 08003, Spain

## Abstract

The biological relevance and dynamics of mRNA modifications have been extensively studied in the past few years, revealing their key roles in major cellular processes, such as cellular differentiation or sex determination. However, whether rRNA modifications are dynamically regulated, and under which conditions, remains largely unclear. Here, we performed a systematic characterization of bacterial rRNA modification dynamics upon exposure to diverse antibiotics using native RNA nanopore sequencing. To identify significant rRNA modification changes, we developed *NanoConsensus*, a novel pipeline that integrates the estimates from multiple RNA modification detection algorithms, predicting differentially modified rRNA sites with very low false positive rates and high replicability. We showed that *NanoConsensus* is robust across RNA modification types, stoichiometries and coverage, and outperforms all individual algorithms tested. Using this approach, we identified multiple rRNA modifications that are lost upon the presence of antibiotics, showing that rRNA modification profiles are altered in an antibiotic-specific manner. We found that significantly altered rRNA modified sites upon antibiotic exposure are located in the vicinity of the A and P-sites of the ribosome, possibly contributing to antibiotic resistance. We then systematically examined whether loss of ‘antibiotic-sensitive’ rRNA modifications may be sufficient to confer antibiotic resistance, finding that depletion of some rRNA modification enzymes guiding dysregulated rRNA modifications confers increased antibiotic resistance. Altogether, our work reveals that rRNA modification profiles can be rapidly altered in response to environmental exposures, and that nanopore sequencing can accurately identify dysregulated rRNA modifications, contributing to the mechanistic dissection of antibiotic resistance. Moreover, we provide a novel, robust workflow to study rRNA modification dynamics in any species using nanopore sequencing in a scalable and reproducible manner.

## INTRODUCTION

Increasing antibiotic resistance among pathogenic bacteria threatens healthcare and the efficacy of the majority of currently known antibiotics^1^. Most clinically used antibiotics inhibit bacterial growth by targeting protein synthesis^2–4^, often through direct binding to the bacterial ribosome, interfering with mRNA translation or blocking the formation of peptide bonds at the peptidyl transferase center. For example, aminoglycoside antibiotics such as kanamycin, streptomycin and neomycin, are potent and broad-spectrum antibacterials that were introduced in the clinic more than five decades ago, which interact with the 16S rRNA at the A site of the ribosome^5^. Unfortunately, their clinical efficacy is seriously threatened by multiple resistance mechanisms^6^. Currently, the most widely disseminated aminoglycoside resistance determinants are drug modification enzymes^7^, but 16S rRNA methyltransferases that modify the drug-binding site have recently emerged as a significant threat that can confer class-wide resistance to these drugs^8, 9^. Thus, detailed studies of these emerging resistance mechanisms are urgently needed.

Bacterial rRNAs contain a large number of methylations that are placed by genomically encoded methyltransferases^10^. While these typically improve ribosome function under most conditions, when challenged with antibiotics the loss of specific modifications can confer low to moderate levels of antibiotic resistance^11–16^ (**Table S1**). For example, loss of the dimethylation of adenine (m^6,6^A) at position 1519 of the 16S rRNA (16S:m^6,6^A1519), which is placed by the genomically encoded methyltransferase *rsmA*, has been shown to confer resistance to kasugamycin^17, 18^, whereas loss of 7-methylguanosine (m^7^G) in the 16S rRNA (16S:m^7^G527), which is placed by the *rsmG* methyltransferase, confers resistance to streptomycin^19^. To better comprehend the role that rRNA methylation dynamics plays in antibiotic resistance, and to decipher whether additional rRNA modifications might be contributing to increased antibiotic resistance, accurate methods to monitor and quantify rRNA modifications are sorely needed^20–22^.

Transcriptome-wide detection of RNA modifications has been typically achieved by coupling either antibody immunoprecipitation or chemical probing with next generation sequencing (NGS) technologies^23–35^. However, limited availability of selective antibodies and/or chemicals only allows for detection of ∼5% of currently known RNA modifications^21, 36, 37^. Moreover, even when these reagents are available, these methodologies have high false positive rates^38^, are often not quantitative^39^ and are inconsistent when using different antibodies^40^, and can only detect one RNA modification type at a time. A promising alternative to NGS-based methods is the direct RNA nanopore sequencing (DRS) platform developed by Oxford Nanopore Technologies (ONT), which can detect diverse types of modified nucleotides in individual native RNA molecules^41–43^. In this platform, RNA molecules are translocated through the nanopores that are embedded in synthetic membranes coupled to an ammeter, causing changes in the ionic current, which are in turn used to identify the underlying nucleotide sequence using machine learning algorithms^44–46^, a process referred to as ‘base-calling’. RNA modifications can then be identified using two main approaches: (i) in the form of systematic base-calling ‘errors’^47–51^, or (ii) in the form of alterations in the current signal (i.e., altered current intensities, dwell times and/or trace) ^43, 52–57^. In recent years, a plethora of algorithms to detect RNA modifications in DRS datasets have been developed^47, 53, 56–59^; however, the overlap between predicted RNA modified sites by different algorithms is poor^49, 57^, limiting our ability to extract meaningful biological conclusions from these datasets. Moreover, it is currently unclear how the performance of each algorithm varies depending on the RNA modification type, modification stoichiometry and sequencing depth, thus limiting the applicability of DRS for the detection of dynamically regulated RNA modifications in biological contexts^45^.

Here, we systematically benchmark diverse RNA modification detection softwares in DRS datasets, across diverse RNA modification types, stoichiometries and sequencing depths. We then propose a novel approach, *NanoConsensus* which uses as input the predictions of diverse RNA modification prediction softwares (EpiNano^47^, Nanopolish^60^, Tombo^56^ and Nanocompore^57^), re-scores them and weights them internally, and finally extracts a robust list of reproducible RNA modification sites that are differentially modified between two conditions (e.g. wild type versus knockout; antibiotic-treated versus untreated). Our results demonstrate that *NanoConsensus* is a robust strategy to detect multiple rRNA modification types simultaneously, outperforming all individual RNA modification softwares tested in this work, detecting RNA modification changes across diverse RNA modification types, with improved sensitivity and specificity across a wide range of modification stoichiometries and sequencing depths.

We then apply *NanoConsensus* to study rRNA modification dynamics on *E. coli* cultures from diverse genetic backgrounds (including strains lacking specific rRNA modification enzymes), which were subjected to either streptomycin (str) or (ii) kasugamycin (ksg) exposure, which bind to the A- and P-site of the ribosome, respectively (**Figure 1A**). We find that upon antibiotic treatment, rRNA modification levels of a subset of sites are significantly decreased, in an antibiotic-dependent manner, and that this loss of rRNA modifications depends on the specific antibiotic employed. Notably, dysregulated rRNA modified sites are spatially located in the vicinity of the A site and the P-site of the ribosome. We show that this loss is not caused by the appearance of mutations in rRNA molecules, nor expression of alternative rRNA operons. Rather, we demonstrate that the loss of rRNA modifications is caused by the *de novo* appearance of a subpopulation of under-modified rRNA molecules that were not present in untreated *E.coli* cultures.

**Figure 1.**
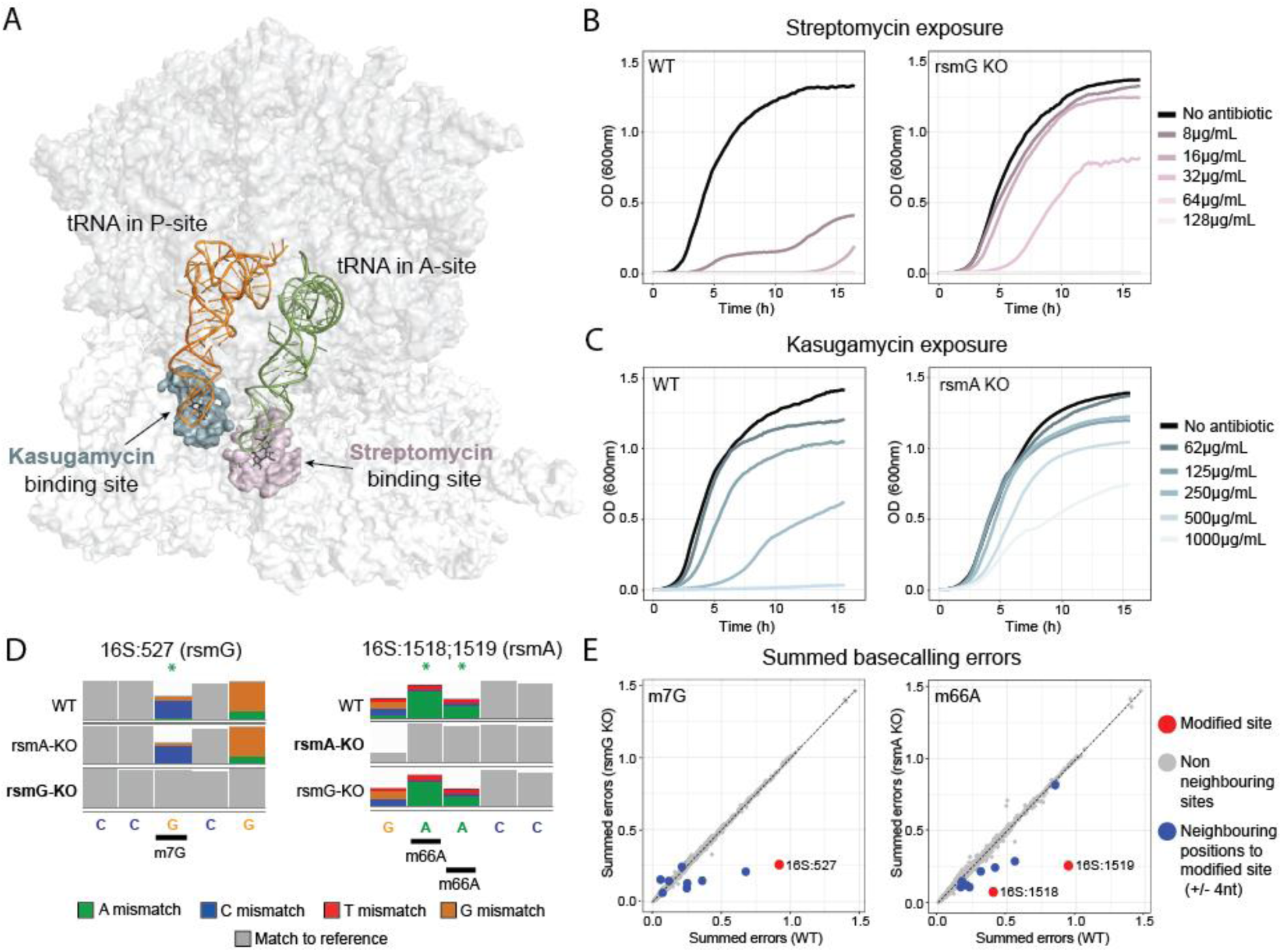
Nanopore direct RNA sequencing can be used to study bacterial rRNA modifications involved in antibiotic resistance mechanisms. **(A)** 3D structure of the ribosome (gray) depicting the tRNA in the P-site (orange) and A-site (green). The residues surrounding the kasugamycin binding site are shown in blue, and are located near the anticodon region of the tRNA that is located at the P-site, whereas residues surrounding the streptomycin binding site are shown in pink, and are located near the anticodon region of the A-site tRNA. Surrounding residues were defined as those that have at least one of its atoms at less than 10Å from the antibiotic. PDB structure corresponds to 7K00. See also *Methods* **(B)** Growth curves of *E. coli* WT (left) and *E.coli rsmG* KO (right) upon increasing concentrations of streptomycin. Data shown represents the average OD_600_ values of 2 independent biological replicates. Each biological replicate was calculated as the median of 3 technical replicates. Data was collected every 9 minutes, during a time course of 16h. **(C)** Growth curves of *E. coli* WT (left) and *E.coli rsmA* KO (right) upon increasing kasugamycin concentrations. Data shown represents the average OD_600_ values of 2 independent biological replicates, and each biological replicate was calculated as the median of 3 technical replicates. Data was collected every 9 minutes, during a time course of 16h. **(D)** IGV snapshots of *E.coli* WT and knockout strains illustrating loss of base-calling errors upon enzyme knockout (*rsmG* and *rsmA*). Positions with mismatch frequencies greater than 0.1 are coloured, whereas positions with mismatch frequencies lower than 0.1 are shown in gray. **(E)** Scatterplots of the summed base-calling error frequencies (sum of insertion, deletion and mismatch frequencies) at each nucleotide position in the knockout strain, relative to WT. The rRNA modified sites that are lost upon knockout of the gene are shown in red (position 0); the neighboring positions (±4 nt) to the rRNA modification site (position 0) are shown in blue. Remaining positions are shown in gray.

Overall, our work reveals that rRNA modifications can be dynamically regulated upon antibiotic exposure, with altered rRNA modification patterns that are antibiotic-specific. Moreover, we demonstrate that *NanoConsensus* is a robust toolkit to study rRNA modification dynamics, that is applicable to diverse RNA modification types, and is robust across varying stoichiometries, with low false positive rates. To facilitate the use and applicability of *NanoConsensus* by future users, we integrate this pipeline into the *MasterOfPores* NextFlow workflow^61^, making the detection of differential RNA modifications DRS datasets simple, traceable and reproducible.

## RESULTS

### Direct RNA nanopore sequencing can identify bacterial rRNA modifications implicated in antibiotic resistance mechanisms

Previous works have shown that bacteria have evolved an effective and elegant way of preventing drug binding to the ribosome by either adding or removing specific rRNA modifications at appropriate sites^62, 63^. Yet, a systematic analysis of how antibiotic exposures affect the stoichiometry and dynamics of rRNA modifications is currently missing. The direct RNA nanopore sequencing (DRS) platform is well-suited to capture the dynamic changes in rRNA modified sites caused by environmental cues, such as antibiotics^43, 64^. However, whether this is the case, and whether nanopore sequencing is sensitive enough to capture rRNA modification changes caused by the presence of antibiotics, remains unknown.

To address this question, we first examined whether *E. coli* mutant strains lacking specific rRNA methyltransferases would show increased antibiotic resistance phenotypes, as reported in the literature. To this end, we cultured *E. coli* strains lacking *rsmG* (responsible for placing 16S:m^7^G527) or *rsmA* (responsible for placing 16S:m^6,6^A1518,A1519) (**Table S1**). Cultures from different genetic backgrounds were then exposed to increasing antibiotic concentrations (streptomycin or kasugamycin, see **Figure 1A**) for which their resistance had been reported, and their growth was monitored during 16 hours. Results confirmed that *rsmG* and *rsmA* knockout strains showed increased resistance to streptomycin and kasugamycin, respectively (**Figure 1B,C**), compared to wild type strains, in agreement with previous literature^17, 19^.

We then examined whether the lack of rRNA modifications previously implicated in antibiotic resistance mechanisms (16S:m^7^G527 and 16S:m^6,6^A1518,A1519) would be identifiable using nanopore sequencing. Previous works studying RNA modifications using nanopore sequencing have mainly focused their efforts on the detection of N6-methyladenosine (m^6^A) ^47, 51, 53, 55, 57, 58, 65^ and pseudouridine (Ψ) ^43, 66–68^ –and to a lesser extent inosine (I)^69^ and 2’-O-methylation (Nm)^43, 70^–, finding that these can be identified with reasonable accuracy. However, whether DRS can detect other less frequent RNA modification types, such as those that are involved in bacterial antibiotic resistance mechanisms, remains unclear. To this end, we sequenced total RNA from *E. coli rsmG*, *rsmA* and *rsmF* (responsible for 16S:m^5^C1407) knockout strains, as well as from the parental *E. coli* wild type strain, using DRS (**Figure 1D**, see also **Figure S1**). *EpiNano-Error*^47^ was used to identify differentially modified rRNA sites by comparing the base-calling ‘errors’ of each knockout strain to those observed in the parental wild type strain, revealing that all 3 modifications examined (m^7^G, m^6,6^A and m^5^C), could be identified in the form of base-calling ‘errors’ ^48, 49^ (**Figure 1D-E**, see also **Figure S1A**). Indeed, we found that base-calling ‘errors’ decreased in the *rsmG*, *rsmA and rsmF* knockout strains (**Figure 1D**, see also **Figure S1B**), supporting that DRS can be used to identify rRNA modification types that are involved in antibiotic resistance mechanisms.

### Poor reproducibility across DRS RNA modification detection algorithms limits our ability to identify antibiotic-induced rRNA modification dynamics

Considering that m^7^G and m^6,6^A-deficient strains (*rsmG* and *rsmA* knockouts) displayed increased resistance to streptomycin and kasugamycin, respectively (**Figure 1B,C**), we then wondered whether the wild type *E. coli* strain would dynamically modulate its rRNA modification levels of 16S:m^7^G527 and 16S:m^6,6^A1518,1519 upon antibiotic exposure. To address this question, we treated wild type *E. coli* cultures with either streptomycin (str) or kasugamycin (ksg) for either 1h or 16h, and sequenced treated and untreated samples using DRS (**Table S2)**. Differences in rRNA modification profiles were determined using *EpiNano-Error*, as well as with 3 additional softwares (*Nanocompore*^57^, *Tombo*^56^ and *Nanopolish*^71^) that employ pairwise comparison approaches to identify differential RNA modifications between two samples, and therefore, are not limited to detecting a single RNA modification type. We should note that *Nanopolish* does not identify RNA modifications *per se*; however, it can be used to ‘resquiggle’ the reads to then predict differences in current intensity at each position^45^.

Our results showed that each algorithm predicted many dynamic sites between the two conditions (antibiotic-treated versus untreated); however, we observed a poor overlap in the predictions made by each algorithm (**Figure S2**), with *Tombo* predicting most of the sites as differentially modified, in agreement with previous works^53, 57^. The poor overlap might be caused by ‘resquiggling’ differences caused by unequal read lengths across the samples ^43, 56^, and/or by unequal coverage along the transcripts. To ensure that unequal coverage along the transcript would not cause biases in the detection of differentially modified sites along the transcript and across algorithms, only full-length reads were kept for downstream analyses (see *Methods* and **Table S2**). In addition, to compare the results obtained by each RNA modification prediction software in a uniform manner, values reported by each software were converted into Z-scores across each transcript, and differentially modified rRNA sites were predicted by comparing the Z-scores at each position to the background median Z-score of each transcript (see *Methods*) (**Figures S3 and S4**).

Using this approach, we found that all 4 softwares showed good overlap of predicted differential sites across biological replicates, within each software (38-69% replicability, depending on the software) (**Figure S5A**). However, we still observed a poor overlap of predicted differentially modified sites across softwares (15-25%) (**Figure S5B**) even when filtering for full-length reads, suggesting that a robust pipeline to detect differentially modified RNA sites from DRS datasets is greatly needed.

### Performance of nanopore-based RNA modification detection algorithms varies depending on the RNA modification type, sequencing coverage and modification stoichiometry

To investigate the reasons behind the poor reproducibility of RNA modification detection across softwares, we systematically analyzed how different features (sequencing coverage, RNA modification type and modification stoichiometry) might impact the performance of each algorithm. To this end, we analyzed DRS datasets from *E. coli* and *S. cerevisiae* wild type and mutant strains lacking distinct RNA modification types at known rRNA positions (see **Table S3**).

Firstly, we examined whether different RNA modification types (Am, m^5^C, m^7^G, m^6,6^A, Um, Ψ) would be detected by each different software. We should note that RNA modifications appear in the form of altered base-calling ‘errors’ and/or current intensities in positions that are neighboring the modified site (i.e. positions −2,−1, +1 and/or +2 relative to the modified site) ^43, 47, 57^. For this reason, we quantified the scores in the 5 nucleotides surrounding the differential modified site (5-mer), and then examined how each modification type was detected by each software (**Figure 2A**). We observed that the position within the 5-mer identified as ‘altered’ was affected both by the RNA modification type and choice of algorithm (**Figure 2A**). For example, *EpiNano* typically identified alterations in the base-calling signatures at the position 0 of the 5-mer, whereas current intensity-based methods, such as *Tombo*, *Nanopolish* or *Nanocompore*, frequently identified the alterations at the neighbouring positions. Moreover, alterations in the signal were often seen at different positions of the 5-mer, even when comparing the same RNA modification type across different algorithms. For example, 2’-O-methyladenosine (Am) could be detected by all 4 softwares, but the altered signal was detected at distinct positions of the 5-mer depending on the algorithm. Altogether, these results suggest that in order to compare the predictions from individual softwares, Z-score information from all positions of the 5-mer should be taken into consideration for downstream analyses, rather than comparing the nucleotide positions predicted by each software, as each software associated the modification signal to distinct nucleotide position(s).

**Figure 2.**
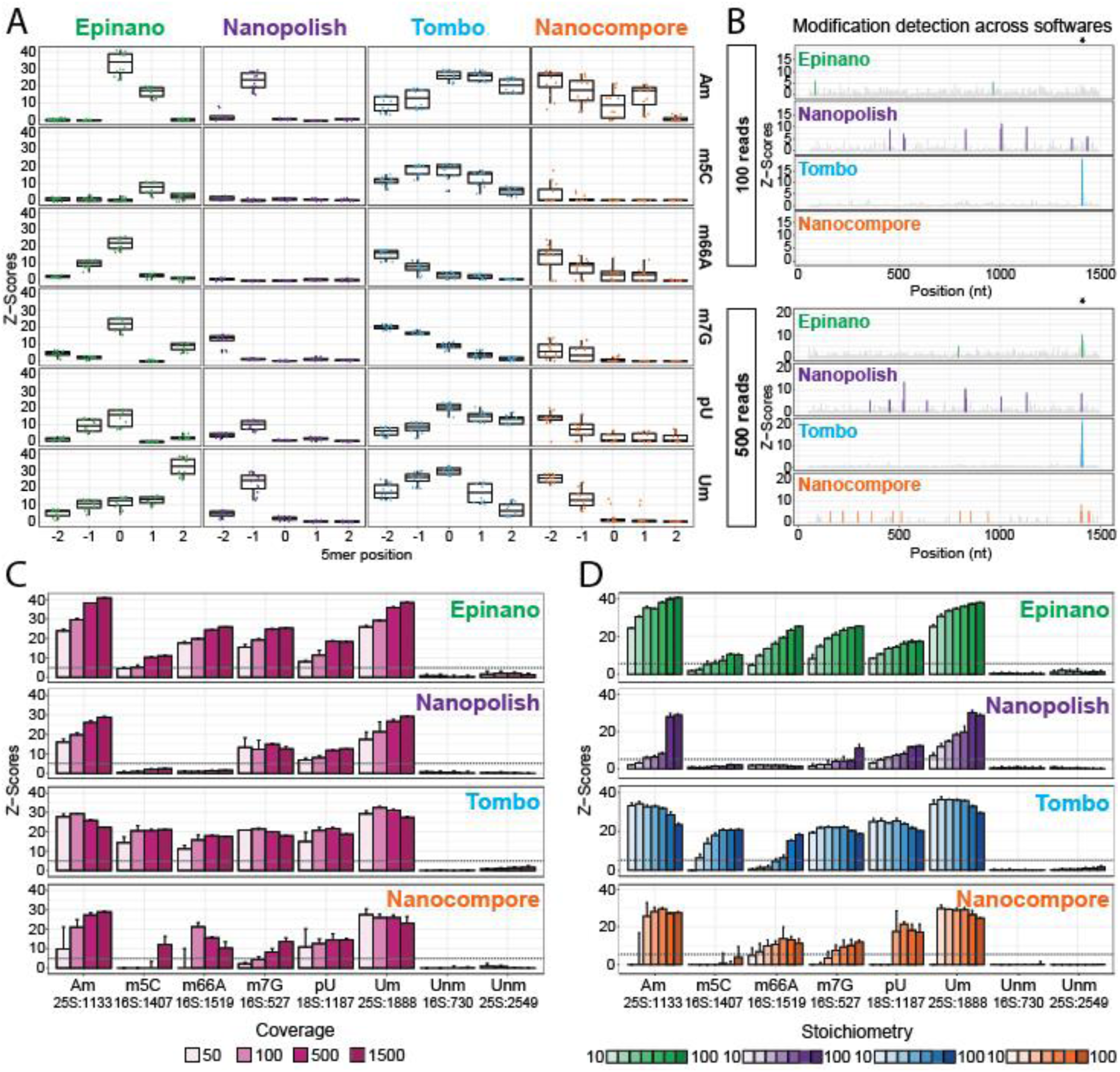
Algorithm performance varies depending on RNA modification type, modification stoichiometry and sequencing coverage. **(A)** Dotplots of Z-scaled differential RNA modification scores along the modified 5- mer obtained by each software when detecting different RNA modification types. In the x-axis, 0 denotes the modified position. Position within the 5-mer of the ‘altered feature’ varies depending on the RNA modification type, but also depending on the software used to identify the modification changes. Box, first to last quartiles; whiskers, 1.5× interquartile range; center line, median. **(B)** Z-scaled RNA modification scores along the *E. coli* 16S rRNA transcript for each individual software tested, obtained when comparing *E. coli* wild type and *rsmF* knockout strains. Analyses have been performed using either 100 (upper panel) or 500 randomly selected reads (lower panel). The modified site that is reported to be lost in the *rsmF* knockout strain (16S:m^5^C1407) is indicated with an asterisk (*). See also **Figures S6-S9**. **(C)** Barplots of median differential modification Z-scores obtained by each software in the 5-mer region centered in each modification type present in the benchmarking. Differential Z-scores are obtained by comparing the wild type and knockout strains, for each given site. Two randomly chosen rRNA modified sites that are not expected to change across any of the datasets have been included in the analyses as controls (16S:730 and 25S:2549). Error bars indicate median ± s.d. of three replicates. Horizontal line shown in gray depicts Z-score = 5. See also **Table S3**. **(D)** Barplots of median Z-scores of the 5-mer regions centered in each modified site analyzed. Samples were analyzed using 1000 reads per sample, and with a diverse range of stoichiometry levels (see *Methods*). Error bars indicate median ± s.d. of three replicates. Horizontal line shown in gray depicts Z-score = 5.

Next, we examined how sequencing coverage affected the performance of each software. To this end, each DRS dataset (**Table S3**) was downsampled into three independent subsets of 50, 100, 500 and 1500 reads (see *Methods*). Each downsampled wild type subset was compared to a downsampled knockout subset of reads, and differential peaks were identified after Z-score normalization (**Figure 2B**, see also **Figures S6-S9**). Our results revealed that the identification of differentially modified sites was strongly dependent on sequencing coverage (**Figure 2C**). Notably, the minimal coverage required to detect a differentially modified site varied both depending on the algorithm as well as on the modification type. For example, loss of m^5^C could be detected by *Epinano* when having a minimum coverage of 500 reads, whereas *Tombo* was able to detect it with a coverage of 50 reads. By contrast, *Nanocompore* was only able to detect this position with a coverage of 1500 reads, and *Nanopolish* was unable to identify this site as differentially modified at the tested coverage conditions. Overall, our results demonstrate that coverage significantly affects the performance of all algorithms, that not all softwares efficiently detect all RNA modification types given a specific sequencing coverage, and that using 50 reads per position (which is the recommended threshold by several of the softwares) is often insufficient to detect RNA modifications, even when the site modified at high stoichiometry.

Finally, we assessed the impact of RNA modification levels (i.e. stoichiometry) in the performance of each algorithm. To this end, we artificially generated samples with decreasing modification stoichiometries (100, 75, 50, 40, 30, 20 and 10%) by mixing reads from wild type and knockout samples at different proportions, with the assumption that wild type samples are 100% modified at the site of interest, and knockout samples are 0% modified (see *Methods*). Subsamplings were performed in triplicate for each RNA modification type and stoichiometry level. We should note that all artificial mixes representing distinct RNA modification stoichiometries contained the same number of reads (n=1000), to ensure that coverage would not be a confounder in the analysis. Our results revealed that stoichiometry had a major effect in the detection of RNA modifications (**Figure 2D**), in agreement with previous works^57, 59^. Notably, we observed that the effect of stoichiometry was strongly dependent on the RNA modification type. For example, Um and Am were detected by most of softwares across all stoichiometries tested; by contrast, the majority of algorithms struggled to detect m^5^C, even at high stoichiometries, with the exception of *Tombo* (**Figure 2D**). Our results showed that *Epinano* and *Tombo* overall performed better than *Nanopolish* and *Nanocompore*, especially at low stoichiometries. We propose that this might be explained by the fact that both use *Nanopolish* algorithm for resquiggling, which has been reported to preferentially resquiggle unmodified reads, resulting in a biased population of resquiggled reads^43^, and consequently, decreased performance in detecting low-stoichiometry RNA modified sites.

### *NanoConsensus* outperforms individual softwares at predicting RNA modifications

Our results showed that RNA modification type, stoichiometry levels and coverage impact the detection of differentially modified sites in DRS datasets (**Figure 2**). Notably, our results also revealed that all softwares examined predicted a significant number of false positives. Indeed, each software identified several ‘differentially modified’ sites when comparing wild type and knockout strains, even though only one RNA modified site is absent in the knockout strain, relative to the wild type (**Figure 2B**, see also **Figures S6-S9**). We reasoned that the detection could be improved if the predictions from different algorithms would be combined in a consensual manner, as this should decrease the number of false positives while retaining the true differentially modified sites. To this end, we developed *NanoConsensus* (**Figure 3A**), an algorithm that reports putative modified regions at per-transcript level, which takes as input the results generated by different RNA modification detection algorithms (*EpiNano*, *Nanopolish*, *Tombo* and *Nanocompore*), and generates as output a list of differentially modified sites that are robustly predicted across softwares and replicates.

**Figure 3.**
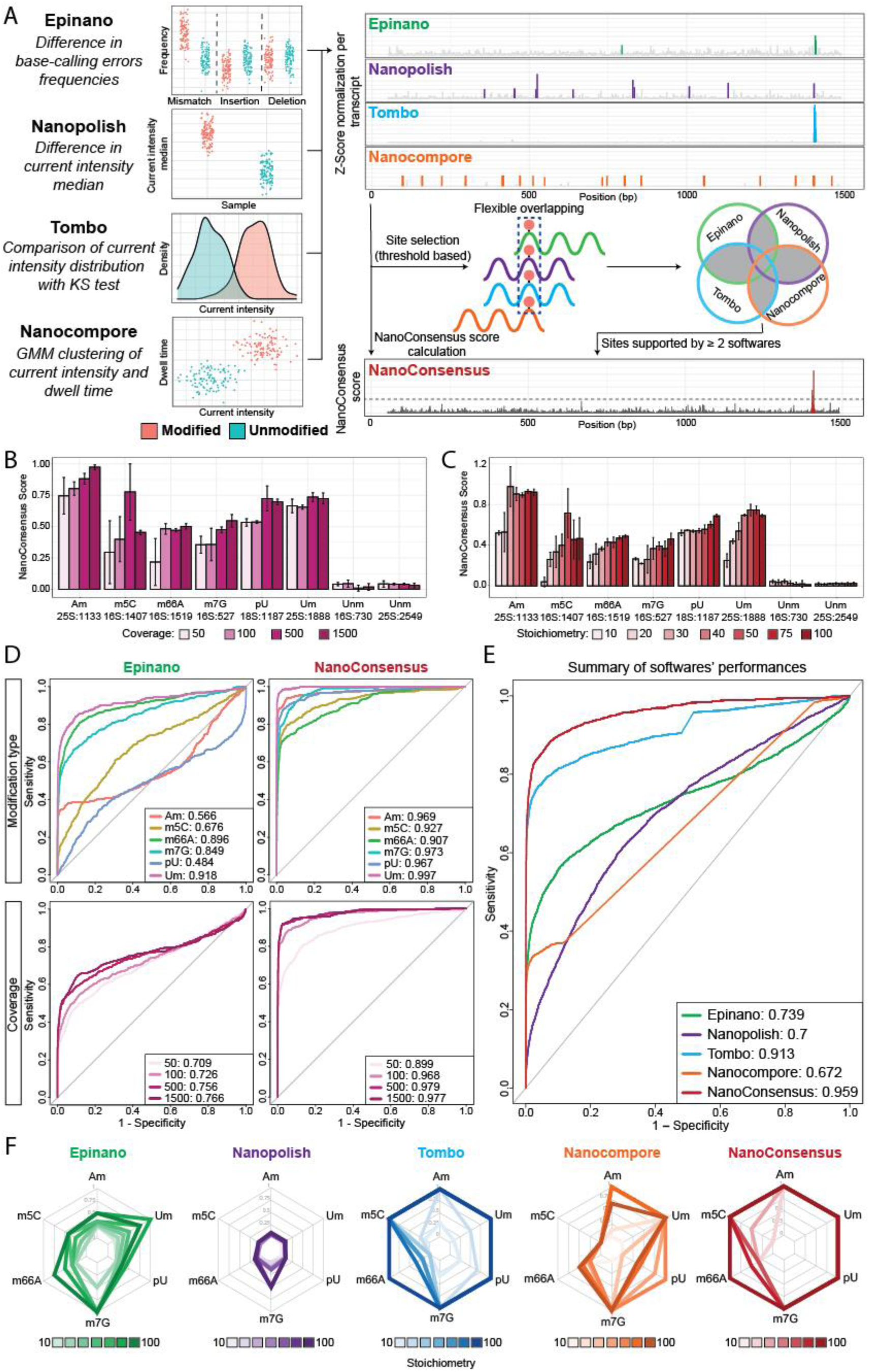
*NanoConsensus* outperforms individual softwares in the prediction of RNA modifications. **(A)** Schematic overview of the *NanoConsensus* pipeline **(B)** Barplot of *NanoConsensus scores* of the 5-mer regions centered in each modified site analyzed. Samples included in the analysis consist of 100% modified samples (WT), with varying read coverage levels, and are the same that were used to generate **FIgure 2C**. Error bars indicate median ± s.d. of three replicates. **(C)** Barplot of median *NanoConsensus scores* of the 5-mer regions centered in each modified site analyzed. Samples were analyzed with a range of stoichiometry levels (and fixed coverage of 1000 reads), which are the same samples that were used to generate **Figure 2D**. Error bars indicate median ± s.d. of three replicates. **(D)** ROC curves from *Epinano* and *NanoConsensus* across a range of modification types and coverage levels. Each curve was computed by merging the data from all stoichiometries and replicates (see *Methods*). At the bottom corner, the legend and AUC values for each ROC curve are shown. See also **Table S5** for median AUC values for all conditions included in the study. **(E)** ROC curves for each software, computed by merging the predictions for all modification types, stoichiometry and coverage levels included in the benchmarking. In the bottom corner, the respective AUC values for each ROC curve are shown. **(F)** Radar plots of positive predictive values (PPV) for each software, RNA modification type and stoichiometry. PPVs were computed using a Z-Score threshold > 5 and a *NanoConsensus* score threshold > 5*(median across transcript). See also **Table S6** for median PPV values.

Briefly, *NanoConsensus* first transforms the predictions from each software into Z-scores, identifies putative modified sites by each software, and assigns a *NanoConsensus score* to each position of the transcript. Because the alteration of the signal caused by a modification does not necessarily overlap across softwares (**Figure 2A**), *NanoConsensus* expands each putative modified site predicted by each software into a 5-mer window, and predicts the overlap of 5-mer regions predicted by each software. Finally, all regions supported by at least two softwares and with a *NanoConsensus* score higher than a threshold (which is dependent on the background ‘noise’, see *Methods*) are reported as ‘differentially modified’ sites (**Figure 3A**).

We then assessed *NanoConsensus’* performance in the detection of RNA modifications across diverse stoichiometry levels, coverage and modification types (**Figure 3B-C**), using as input the same datasets used to assess the performance of each individual software (**Figure 2**, see also *Methods*). To compare the performance of *NanoConsensus* to that of each individual software, Receiver Operating Characteristic (ROC) curves were built for each algorithm, modification type, coverage and stoichiometry, and the comparative performance was assessed by comparing the Area Under the Curve (AUC) values for each ROC curve. Our results showed that *NanoConsensus* was more robust in predicting RNA modifications across diverse modification types, stoichiometries and coverage (**Figure 3D**, see also **Figures S10 and S11**). Indeed, *NanoConsensus* globally outperformed all individual softwares in terms of AUC values when combining all the subsets (RNA modification types, stoichiometries, and coverage) included in the benchmarking (**Figure 3E**).

Finally, we examined the number of false positives reported by each individual software, compared to *NanoConsensus*. To this end, we computed the positive predictive values (PPV) across diverse stoichiometries and RNA modification types, for each algorithm. The PPV reflects the proportion of true positives (TP) among the set of predicted ‘positive’ (P) differentially modified sites. Our results showed that the PPV was typically lower with the decrease in modification stoichiometry and coverage, and that it was also dependent on the RNA modification type (**Figure 3F**, see also **Figure S12**). Globally, we found that *Nanopolish* showed the worst performance in terms of PPV, followed by *EpiNano* and *Nanocompore*. On the other hand, *NanoConsensus* showed the best performance in terms of PPV across all stoichiometries and modification types, implying that it reports fewer false positives, compared to individual softwares tested.

### Aminoglycoside exposure leads to changes in the modification pattern of the 16S in the bacterial ribosome

We then used *NanoConsensus* to re-examine the question of how antibiotic exposures, such as kasugamycin or streptomycin, may impact the rRNA modification patterns of the bacterial ribosome. To this end, *E. coli* cultures were grown in LB until log phase, and were then exposed to streptomycin (str), kasugamycin (ksg) or left untreated (**Figure 4A**). Cultures were collected 1h or 16h post-treatment, and the experiment was performed in independent biological replicates, on different days. DRS libraries from total RNA were built as previously described ^43^ (see *Methods*), and the data was analyzed in a pairwise manner using *NanoConsensus*. This analysis revealed multiple regions (5 in 16s rRNA, 1 in 23s rRNA) that were ‘differentially modified’ in both replicates (**Figure 4B**, labeled in red). Notably, all the identified regions located in the 16S rRNA overlapped with known rRNA-modified sites (**Table S7**). In terms of directionality of the RNA modification change, all sites identified as differentially modified decreased in their modification stoichiometry upon antibiotic exposure (**Figure S13**). We should note that increased duration of antibiotic treatment (1h or 16h) (**Figure 4B**); did not significantly change the set of differentially modified sites identified in the case of kasugamycin, whereas in the case of streptomycin, some differentially rRNA modified sites were lost with extended exposures.

**Figure 4.**
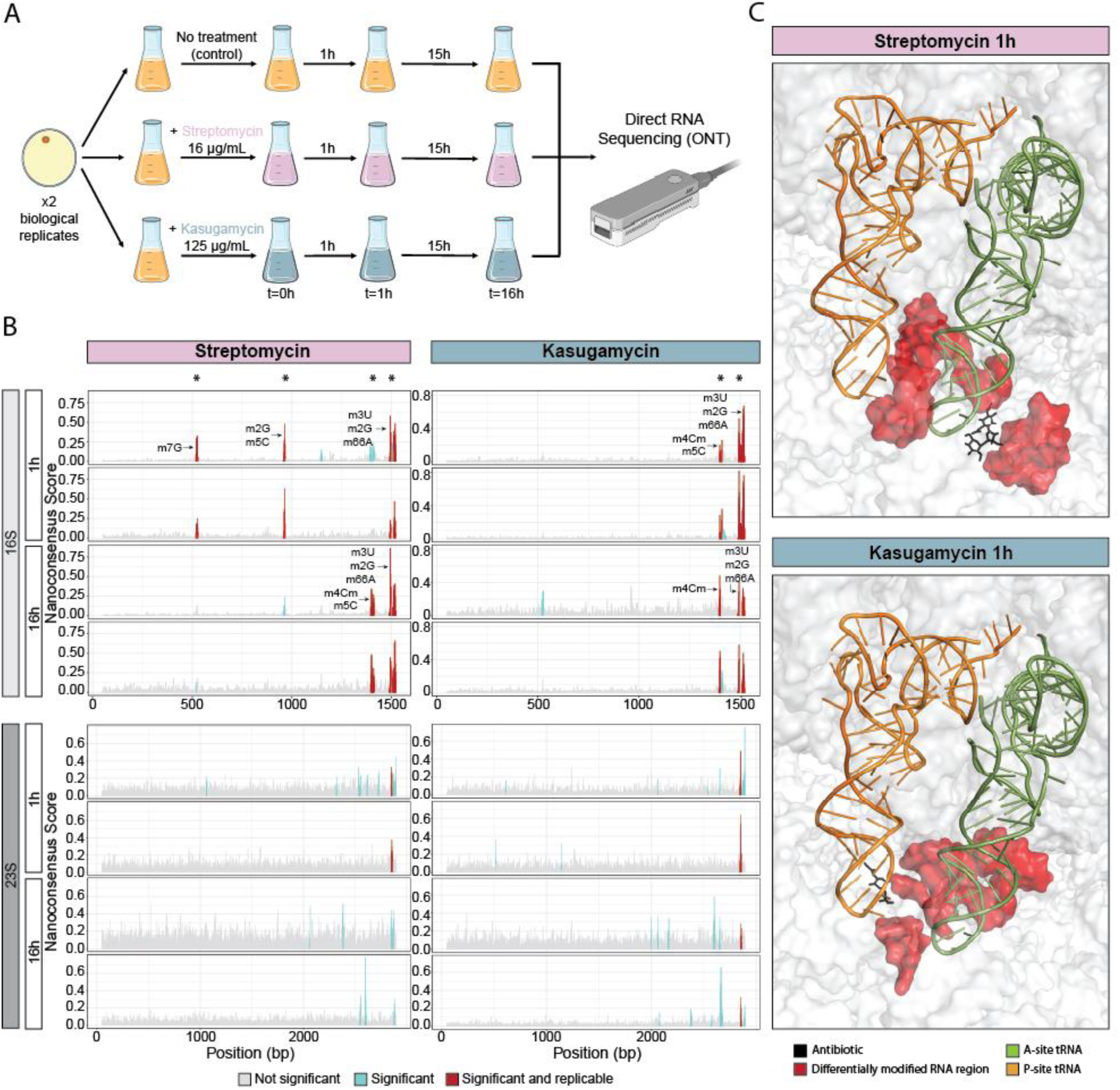
Direct RNA sequencing results of *E. coli* samples exposed to streptomycin and kasugamycin. **(A)** Experimental design of the exposure of non-resistant E.coli to both streptomycin and kasugamycin. **(B)** *NanoConsensus* scores along the 16S rRNA (upper panel) and 23S rRNA (lower panel) of *E. coli* samples collected after 1h or 16h exposure to streptomycin (left panel) or kasugamycin (right panel). In grey, non-significant positions; in blue, regions identified by *NanoConsensus* in only one replicate; in red, regions identified in both replicates. Known modified sites found in replicable differentially modified regions identified by *Nanoconsensus* are also shown. **(C)** 3D structure of the bacterial ribosome (PDB: 7K00), highlighting the differentially modified rRNA residues upon streptomycin (upper panel) or kasugamycin (lower panel) exposure identified in this work. Differential rRNA modified residues are shown in red; in black, the antibiotic; in light green and orange, the tRNA molecule located in the A and P-sites, respectively.

Finally, we examined the 3D location of ‘differentially modified’ regions upon antibiotic treatment within the ribosome structure, and found that these regions were very close in the 3D space despite being far in the linear rRNA molecule (**Figure 4B**). More specifically, these regions were all located within the A- and P-sites of the ribosome (**Figure 4C** see also **Table S7**), thus matching the binding site areas of streptomycin and kasugamycin, respectively (**Figure 4C**). Thus, our results suggest that the alteration of a subset of rRNA modification might constitute an adaptive response of bacterial cells to diminish the binding affinity of kasugamycin and streptomycin in the P-site and A-site, respectively. Altogether, our results demonstrate that rRNA modification levels can be dynamically regulated upon environmental exposures, and that rRNA modification dysregulation might constitute a common molecular mechanism employed by bacterial species to increase their tolerance towards antibiotics binding the ribosome.

### Mutations or differential rRNA usage do not explain the observed differences in DRS profiles

We and others have shown that differential RNA modifications in DRS datasets cause alterations in the current intensities and/or base-calling ‘error’ patterns ^45, 73^. However, changes in the nucleotide sequence (e.g. SNPs, alternative rRNA usage) can also lead to such alterations, and thus could constitute a confounder in the analysis, misguiding the detection of differential rRNA modifications.

To exclude such possibility, we examined whether the use of alternative *E. coli* rRNA genes could explain our results. *E. coli* has seven rRNA operons encoded in its genome, with small sequence differences among them^74, 75^. Notably, previous works have shown that stress can lead to change in its operon usage^76, 77^. To examine whether differential rRNA operon usage could explain our observations, we performed a multiple sequence alignment of the 7 annotated *E. coli* 16S rRNA sequences, finding that all 7 rRNA sequences were identical across operons within regions identified as ‘differentially modified’, (**Figure 5A**) demonstrating that differential rRNA operon usage could not explain the observed changes in the DRS data.

**Figure 5.**
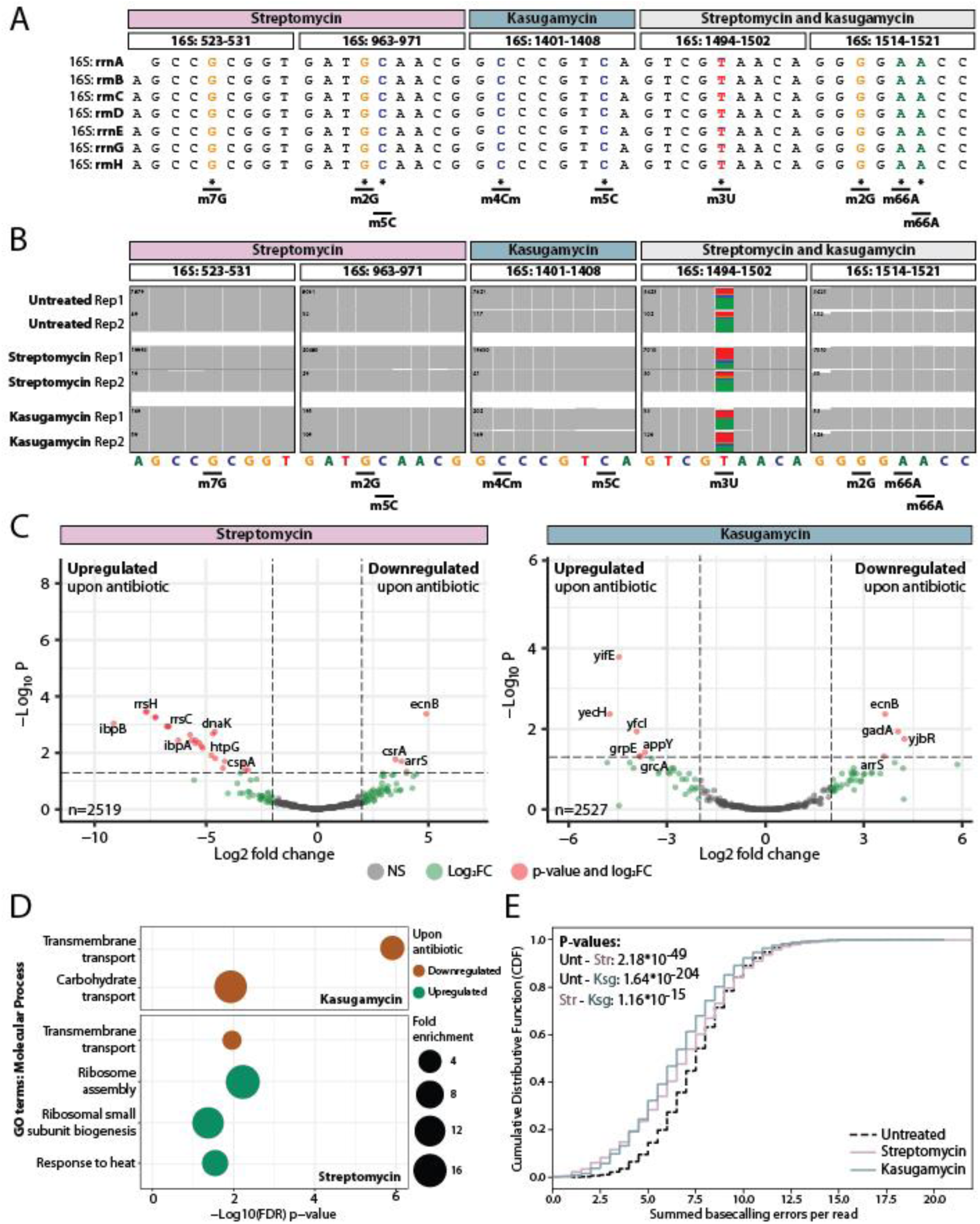
Streptomycin exposure leads to increased *de novo* synthesis of rRNA molecules, some of which lack a subset of rRNA modifications. **(A)** Multiple sequence alignment (MSA) of the seven rRNA operons sequences in all regions identified by NanoConsensus upon antibiotic exposure. **(B)** IGV tracks of nanopore cDNA reads aligning to the ribosome sequence in all regions identified by *NanoConsensus* upon antibiotic exposure. Positions with mismatch frequencies greater than 0.1 are coloured, whereas positions with mismatch frequencies lower than 0.1 are shown in gray. **(C)** Differential expression analysis results between treated and untreated samples. In gray, non-significant genes; in green, genes with abs(log2FC) > 2 and; in red, genes with abs(log2FC) > 2 and p adjusted value >=0.05. **(D)** GO term enrichment analysis using molecular process terms using as input all genes with abs(log2FC) > 2, which corresponded to n=195 (upregulated) and n=214 (downregulated) genes upon streptomycin treatment, and n=197 (upregulated) and n=203 (downregulated) genes upon kasugamycin treatment. In orange, terms related to downregulated genes upon antibiotic exposure and; in green, related to upregulated genes. **(E)** Cumulative distribution of summed basecalling errors at per read level from both untreated (n=10,953 reads), str-treated (n=3,689 reads) and ksg-treated (n=14,376 reads) samples. In the upper-left corner, p-values between distributions calculated with the KS test are shown.

Another possible confounder in the rRNA modification analysis could be the presence of mutations that might arise at the genomic level upon bacterial divisions, which might be selected for if they confer a selective advantage upon antibiotic exposure^78^. To examine whether rRNA mutations could explain our observations, we sequenced antibiotic-treated and untreated samples using Nano3P-seq^79^, a cDNA-based nanopore sequencing method that efficiently captures both the coding and non-coding transcriptome, regardless of their tail composition (see *Methods*). Analysis of rRNA molecules sequencing using Nano3P-seq revealed no mutations at the rRNA regions identified as “differentially modified” by *Nanoconsensus* (**Figure 5B**). Altogether, our analyses support that the changes seen in the DRS data are caused by alterations in the rRNA modification patterns, and not by changes occuring at the rRNA nucleotide sequence or caused by alternative use of rRNA operons.

### De novo rRNA transcription occurs upon streptomycin exposure

Our results show that alterations in bacterial rRNA modification patterns can already be seen 1h after antibiotic exposure (**Figure 5A-B**), demonstrating that rRNA modifications can be rapidly regulated upon environmental exposures. This, in turn, opens new questions regarding how these differentially modified rRNA molecules are produced.

To decipher the mechanism(s) used by bacterial cells to alter their rRNA modification profiles, we first examined whether specific pathways might be altered upon antibiotic exposure. To this end, we sequenced ribodepleted RNA samples from untreated, ksg-treated and str-treated cultures (1h post-antibiotic exposure) using Nano3P-seq, in biological duplicates. We performed differential expression analysis (**Figure 5C** and **Table S8,** see also *Methods*) and GO term enrichment analysis on the set of genes that had at least 4-fold changes in their expression levels (**Figure 5D**), which revealed that both upon kasugamycin and streptomycin treatment, transmembrane transport-associated genes are downregulated, possibly to reduce the antibiotic uptake. In addition, we also found that *de-novo* ribosome synthesis genes were significantly upregulated upon streptomycin treatment, suggesting that the origin of lowly modified 16S rRNA molecules might arise from newly synthesized 16S rRNA molecules.

To test this hypothesis, we examined the cumulative distribution of per-read ‘errors’ (which can be used as a proxy for the relative number of RNA modifications per molecule) from str-treated, ksg-treated and untreated full-length DRS reads (see *Methods*). Our analysis revealed that a population of lowly modified reads (i.e., summed errors < 4) was present in both str- and ksg-treated samples, but was largely absent in untreated samples, suggesting that lowly-modified rRNA reads appear in response to antibiotic treatments (**Figure 5E**). Similar results were obtained using PCA analysis of per-read rRNA modification profiles, which also showed that most of the lowly-modified rRNA reads were mainly appearing upon antibiotic exposure (**Figure S14**).

### Lack of individual methyltransferases does not lead to an increased resistance to streptomycin nor kasugamycin

Previous works have reported that the lack of *rsmA* (responsible of placing m^6,6^A in 16S:1518-1519), and *rsmG* (responsible of m^7^G in 16S:527) leads to increased resistance to kasugamycin and streptomycin, respectively. In our work, we find that modification levels of both 16S:m^6,6^A1518-1519 and 16S:m^7^G527 are significantly decreased upon antibiotic exposure, suggesting an adaptive response of bacteria in their rRNA modification profiles to increase their resistance to antibiotics. Notably, we also found that additional rRNA modified sites, apart from these two, were significantly decreased in their modification levels upon antibiotic exposure (**Figure 4B**). Therefore, we wondered whether the lack of methyltransferases responsible for placing rRNA modifications that we find dysregulated upon antibiotic treatment, in addition to *rsmA* and *rsmG*, might lead to increased antibiotic resistance.

To this end, we cultured seven *E. coli* knock-out strains (listed in **Table S1**), including *rsmA* and *rsmG*, that we identified as responsible for placing rRNA modifications that are differentially methylated upon streptomycin and/or kasugamycin exposure (**Figure 4B**). Each strain was cultured under a broad range of antibiotic (kasugamycin or streptomycin) concentrations, and their growth was monitored for 16 hours (**Figure 6A** and **S15**). To assess whether the lack of a given methyltransferase enzyme led to increased antibiotic resistance, we compared the growth of each knockout to that of the wild type strain by subtracting the AUC values of the growth curves of each strain and antibiotic concentration relative to the same conditions in the wild type strain. Our results showed that the lack of *rsmA* and *rsmG* led to increased antibiotic resistance (**Figure 6B**), as expected. However, the lack of other methyltransferases examined did not lead to increased antibiotic resistance, despite being responsible for placing rRNA modifications at sites that we find differentially modified upon antibiotic exposure. The possible synergistic effect of removing several of these enzymes simultaneously remains unknown.

**Figure 6.**
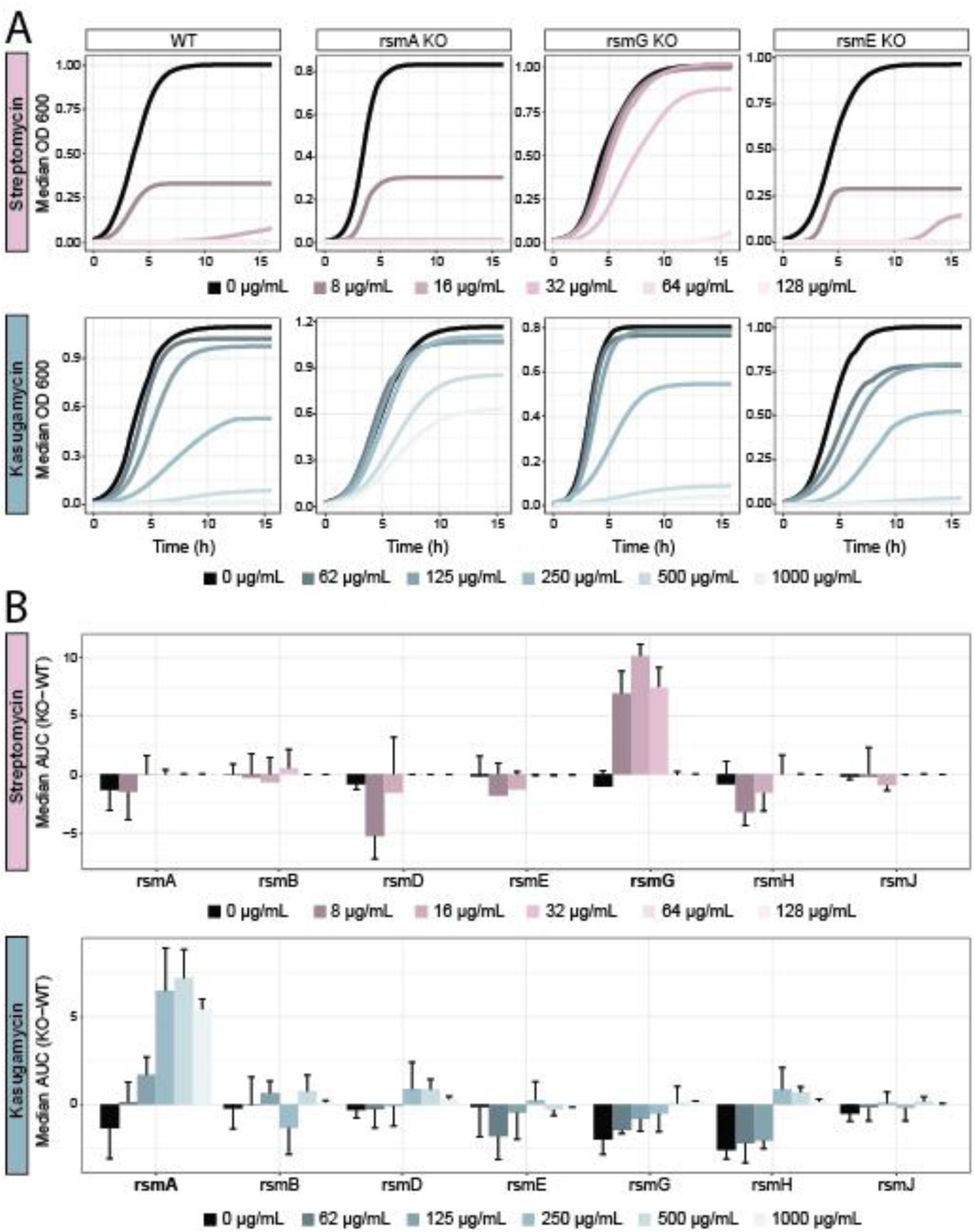
Depletion of *rsmA* and *rsmG*, but not other methyltransferases, leads to increased resistance to kasugamycin and streptomycin. **(A)** Median growth curves from four biological replicates of different *E. coli* strains (wild type, *rsmA* knockout, *rsmG* knockout and *rsmE* knockout) across a range of antibiotic (streptomycin, upper panels; kasugamycin, lower panels) concentrations. Data for each biological replicate was computed as the median of three technical replicates. Growth curve data was fitted to a logistic curve. See also **Figure S15** for growth curves in additional *E. coli* strains tested (*rsmB*, *rsmD*, *rsmH* and *rsmJ* knockout strains). **(B)** Barplots depicting Median Area Under the Curve (AUC) difference in growth curves between wild type and knockout strains (KO-WT), for every KO strain and across all antibiotic concentrations tested. In bold, the strains that are reported to have an increased resistance to each specific antibiotic. Error bars indicate standard deviation of four biological replicates.

## DISCUSSION

A plethora of computational methods for detecting RNA modifications in nanopore direct RNA sequencing (DRS) datasets have been developed in recent years^43, 48–50, 52–58^. These methods typically predict RNA modifications either through the analysis of the raw signal features (e.g., current intensity and/or dwell time) ^43, 52–58^ or in the form of differential base-calling ‘errors’^47–51^. Previous works have compared the performance of some softwares in detecting m^6^A modifications in DRS datasets, finding a poor overlap between the predictions across softwares^53, 57, 58^. However, their comparative performance in detecting other RNA modification types (e.g., Ψ, Nm, ac^4^C) has so far not been assessed. Thus, it is unclear which software(s) should be used to study rRNA modification dynamics.

To tackle this question, here we benchmarked four softwares that can in principle detect diverse RNA modification types in DRS datasets, and assessed their ability in detecting diverse RNA modification types (i.e., Ψ, Am, Um, m^6,6^A, m^7^G and m^5^C) across diverse stoichiometries and sequencing coverage ranges. Our results showed that softwares relying on signal intensity features were better than those using basecalling errors at detecting some RNA modification types (e.g., m^5^C), but the opposite was seen for other RNA modification types (e.g., m^6,6^A). Notably, we also found that in cases where RNA modifications showed a ‘spread of signal’ along several nucleotides, algorithms often identified different positions as the ‘differentially modified site’, partly explaining the poor overlap of RNA modification predictions across softwares (**Figure S2**). In addition, our analyses showed that both coverage and stoichiometry levels had a strong impact in the performance of the algorithms (**Figure 2B-D**), lowering both their sensitivity and specificity (**Figures S6-S10**). To overcome these limitations, we designed and implemented *NanoConsensus*, a workflow that identifies differentially modified RNA sites between two samples using as input the predictions from 4 different softwares, allowing for sliding-window overlaps to identify common predictions across softwares. We found that *NanoConsensus* outperformed all 4 individual softwares, and was robust across a range of RNA modification types, stoichiometry and coverage levels (**Figure 3D-F**). To facilitate the use of this tool, we have made *Nanoconsensus* available as Nextflow module ^80^, and have integrated it within the MasterOfPores ^61^ Nextflow workflow, to ensure reproducibility, scalability and traceability of the analyses performed. We should note that ‘consensus’ approaches have also been successfully employed to improve the detection of DNA modifications in nanopore sequencing data ^81^.

Once the performance of *Nanoconsensus* was benchmarked and optimized, we applied it to investigate whether bacterial rRNA modification profiles were altered upon antibiotic exposure, and determine whether variation in rRNA modification levels might constitute a natural mechanism that bacteria use to increase their resistance upon antibiotic exposure. To this end, we sequenced total RNA from streptomycin-treated, kasugamycin-treated and untreated *E. coli* cultures using nanopore DRS, and compared the rRNA modification profiles of treated and untreated cultures. We found that antibiotic exposure significantly altered the 16S rRNA modification profiles of the bacterial ribosome (**Figure 4B**). Notably, differentially modified rRNA sites were located within the A and P-sites (**Figure 4C** and **Table S7**), which correspond to the binding regions of the streptomycin and kasugamycin, respectively. The fact that rRNA modifications are altered precisely in the vicinity of the antibiotic binding site suggests that this phenomenon is a specialized response from bacteria to the specific antibiotic in question, possibly as a means to decrease the binding affinity of the antibiotic to the bacterial ribosome.

Previous works using DRS have reported differential RNA modification patterns upon heat and oxidative stress in snRNAs, snoRNAs and mRNAs, but not in rRNAs ^43, 64^, suggesting that rRNA modifications might not be dynamically regulated upon environmental conditions. Contrary to previous observations, our work demonstrates that rRNA modifications can indeed be dynamically regulated upon environmental conditions, at least in the case of antibiotic exposure(s). This observation, in turn, opens new questions regarding what might be the mechanism(s) that bacteria use to achieve the outcome of differential rRNA modification. To answer this question, we performed single molecule analysis of rRNA modification patterns (**Figure 5E** and **Figure S14**) as well as differential expression analysis of treated and untreated cultures using Nano3P-seq (**Figure 5C**). These efforts revealed that antibiotic-exposed bacteria have increased *de novo* expression of under-modified rRNA molecules, which ultimately leads to significantly altered rRNA modification profiles. It remains unclear, however, how antibiotics lead to altered rRNA maturation processes and to the production of under-modified rRNA molecules.

Various resistance mechanisms, including enzymatic detoxification, target alteration (rRNAs and ribosomal proteins) and reduced accumulation (impermeability and efflux) have been shown to be involved in bacterial resistance to protein synthesis inhibitors^82^. Notably, rRNA alteration has recently gained increased interest due to several recent studies showing that some chloramphenicol-florfenicol resistant (Cfr) bacteria were carrying a gene encoding for a mutant version of an rRNA methyltransferase^83, 84^. Here we systematically examined whether the loss of individual methyltransferases, responsible for placing rRNA modifications that we identified as differentially modified upon antibiotic exposure (**Table S7**), might alter the resistance of bacteria towards the antibiotic in question. We found that only the loss of *rsmG* and *rsmA,* but not other methyltransferase enzymes examined, led to a resistance phenotype (**Figure 6A-B**, see also **Figure S15**). It is possible that the lack of these additional enzymes is not *per se* enough to lead to increased resistance, but may have a synergistic effect when lost simultaneously with *rsmG* and/or *rsmA*. In this regard, recent studies have reported that some *Mycobacterium tuberculosis* antibiotic-resistant strains carry multiple mutations in several methyltransferase genes ^85^ that are ortholog to the genes identified by *Nanoconsensus*, suggesting that the lack of these additional methyltransferases might constitute a selective advantage towards antibiotic resistance.

While our work establishes a robust framework to identify differential rRNA modification patterns across conditions using DRS (**Figure 2**), we should note that *Nanoconsensus* has some limitations, as well as ample room for future improvement. Firstly, it does not provide information regarding the directionality (i.e., increase or decrease) of the rRNA modification changes between two conditions, although this limitation can be partly alleviated by using the *Epinano* results that are generated as part of the *Nanoconsensus* pipeline (see **Figure S13**). Secondly, *Nanoconsensus* lacks single nucleotide resolution; consequently, if several RNA modifications are located within the differentially modified regions identified, it is not possible to determine which specific rRNA modification is changing. Despite these limitations, we demonstrate that *Nanoconsensus* can be used to reveal antibiotic-dependent rRNA modification changes, contributing to the understanding of how bacteria tune their ribosomes as a response to environmental conditions. Notably, this knowledge may be applied in the future to improve the design of antibiotic chemical structures such that they have improved binding efficiencies towards both the fully-modified and under-modified ribosomal A and P-site regions.

## MATERIALS AND METHODS

### Bacterial strains and culturing

*E. coli* strains used in this study (*rsmA*:JW0050, *rsmG*:JW3718 and *rsmF*:JW5301, see **Table S1** for additional details) were obtained from the Keio Knockout Collection ^86^, including the reference wild type strain (BW25113). Knockout strains have a kanamycin cassette replacing the depleted gene. Strains were plated in LB-agar plates (*E. coli* BW25113) or in LB-agar plates supplemented with 25 μg/mL kanamycin (in the case of *E. coli* knockout strains).

### Growth curves of antibiotic-resistant *E. coli* strains

For each strain, *E. coli* starter cultures were grown O/N in 4 mL tubes at 37°C and 180 rpm. Aliquots from the overnight cultures were added into a final volume of 200μl in 96-well plates with an initial OD_600_ of 0.01. Different antibiotic concentrations (streptomycin: 8,16, 32, 64 and 128μl/mL; kasugamycin: 62, 125, 250, 500 and 1000μl/mL) were tested in both non-resistant and resistant strains. *E.coli rsmF* KO strain was not tested for kanamycin resistance as its KO was generated through the insertion of a kanamycin resistance cassette ^86^. Bacteria was grown at 37°C and OD_600_ was measured every 10 minutes with a TECAN M200 Plate Reader. We should note that the *rsmF* knockout strain was not included in the growth curve experiments as this mutant is expected to show increased resistance to kanamycin, and all Keio collection mutants contain a kanamycin resistance cassette replacing the gene that has been removed ^86^. All experiments were performed in biological replicates (n=4) for each strain and condition, and each strain and condition was measured at least in two independent plates and in two different days.

### Bacterial exposure to antibiotics

*E. coli* BW25113 starter cultures were grown O/N in 4mL of LB at 37°C and 180 rpm. Three 500mL LB cultures were then inoculated with the starter culture, to an initial OD_600_ ≈ 0.01. OD_600_ was monitored every hour with a spectrophotometer until all cultures reached OD_600_ ≈ 0.4 (early log phase). Then, the first 500mL culture was supplemented with streptomycin 10mg/mL (final concentration = 16μl/mL) (Sigma-Aldrich, #S6501-5G), the second 500mL culture was supplemented with kasugamycin 40mg/mL (final concentration = 250μl/mL) (Merck, #32354-100MG) and the third culture was supplemented with the equivalent volume of water (no antibiotic). OD_600_ was measured at time points 0h, 1h and 16h after antibiotic addition (or no antibiotic). 50mL from each erlenmeyer were taken at each of the three time points, and were immediately centrifuged in a pre-chilled benchtop centrifuge at 16,000g 4°C for 10 minutes. Supernatant was discarded, and pellets were stored at −80°C until further use. This experiment was repeated with three independent biological replicates, which were cultured in different days.

### *E. coli* total RNA extraction

Frozen pellets were thawed on ice, followed by the addition of 400uL of Trizol (Life Technologies, #15596018). After five minutes incubation at room temperature, 90uL of chloroform (Vidra Foc, #C2432) was added and mixed thoroughly by inversion. Samples were centrifuged for 15 minutes at 16,000g at 4°C. Supernatant was kept and an equal volume of 70% ethanol was added in a new eppendorf tube. Afterwards, small RNAs were removed with the Qiagen RNeasy Mini Kit following manufacturer’s recommendations (Qiagen, #74104). Samples were DNAse-treated using Turbo DNAse (Life Technologies, #AM2239) for 10 minutes at 37°C followed by a final clean-up with Qiagen RNeasy MinElute Kit (Qiagen, #74204) using default manufacturer’s protocol (which keeps RNAs > 200nt) to remove excess of small RNAs that are typically present in *E. coli* total RNA extracts. RNA integrity was assessed using TapeStation (**Figure S16**) and quantified using Nanodrop.

### Direct RNA nanopore library preparation and sequencing

DNAse-treated *E.coli* total RNA (600 ng per sample) was *in vitro* polyadenylated using *E. coli* Poly(A) polymerase (New England Biolabs, #M0276L) using manufacturer’s recommendations with minor changes (the polyA tailing reaction was carried out for 10 minutes at 37°C, instead of 30). Poly(A)-tailed total RNA was then prepared for sequencing using the direct RNA sequencing kit (SQK-RNA002), following the protocol guidelines (version: *DRCE_9079_v2_revM_14Aug2019*), with minor changes: RNA was linearized using SuperScript IV (Thermo Fisher Scientific, #18090010) and incubated for 15 minutes at 55°C followed by a heat inactivation step (80°C for 10 min), or linearized using Maxima enzyme (Life Technologies, #EP0751) and incubated for 30 minutes at 60°C followed by a heat inactivation step of 5 minutes at 85°C (see **Tables S2 and S3**).

### Base-calling, demultiplexing and mapping of yeast and bacterial direct RNA sequencing runs

Raw fast5 reads from yeast and bacterial samples (**Tables S2 and S3**) were analyzed using the *MasterOfPores* (MoP) version 2 Nextflow workflow ^61^. Briefly, the *mop_preprocess* module was used to demultiplex the FAST5 reads using DeePlexiCon with default parameters ^87^. Demuxed FAST5 were then base-called using *Guppy* 3.1.5 (Oxford Nanopore Technologies) with the model *rna_r9.4.1_70bps_hac*, and aligned using graphmap v 0.5.2 with *-v 1 -K fastq* parameters to the the *E. coli* rRNA reference transcriptome, which comprised the 5S rRNA, the 16S rRNA and the 23S rRNA sequences (available in Github: *NanoConsensus/ref/Escherichia_coli.rRNA.fa*).

### Detection of RNA modifications using diverse algorithms and implementation into the MasterOfPores Nextflow workflow

To detect differential RNA modifications across strains, each control condition (e.g. *E. coli* wild type BW25113) was compared in a pairwise manner to each of the knockout strains (e.g. to the *E. coli* knockout strains JW5301, JW0050 and JW3718, respectively) (**Table S1**). For each pairwise comparison, four RNA modification detection algorithms were run in parallel: i) EpiNano ^49^, ii) Nanopolish ^71^, iii) Tombo ^56^ and iv) Nanocompore ^57^. EpiNano (version 1.1.1) was run with default parameters to extract per-site and per-kmer base-calling errors from every sample. Nanopolish (version 0.11.1) was used to resquiggle the basecalled fast5 using *nanopolish eventalign* (*--samples --print-read-names --scale-events*). Then, current intensity per position of the resquiggled reads and the coverage was extracted with a in-house python script, available in GitHub (https://github.com/biocorecrg/MOP2/blob/main/mop_mod/bin/mean_per_pos_v3.py) ^61^. Tombo (v1.5) was first used to resquiggle the reads using *tombo resquiggle* (*-rna* option). The function *tombo detect_modifications level_sample_compare* with *--store-p-value* was then used to perform RNA modification detection. Finally, p-values at position level together with coverage data were retrieved with *tombo text_output browser_files.* Nanocompore (version 1.0.0) was run through the command *nanocompore sampcomp ( --sequence_context 2 --downsample_high_coverage 10000 --pvalue_thr 1 --logit --comparison_methods GMM,KS,MW,TT)* (**Figure S17**).

### Improving the speed and reducing computational and time requirements of module *mop_mod from MasterOfPores Nextflow workflow*

Nanopolish eventalign output results are stored in csv files, which we found were very slow to parse, increasing the computational time required to perform all the downstream analyses, thus limiting its applicability transcriptome-wide. To increase the speed of the analysis, two in-house scripts based on the *Apache Parquet* format ^88^ were implemented in Python3 (available in GitHub: https://github.com/biocorecrg/MoP3/mop_mod/bin/mean_per_pos.py and https://github.com/biocorecrg/MoP3/mop_mod/bin/Merging_processed_nanopolish_data.py). On the other hand, *Tombo* output files are given in the form of wig and bedgraph files. Therefore, to increase the efficiency of file parsing, we converted Tombo output files into bigWig files and processed them at per-transcript level using the pyBigWig package ^89^. These scripts have been implemented in the MasterOfPores version 2.0 NextFlow workflow ^80^, which is available in GitHub (https://github.com/biocorecrg/MoP2).

### Prediction of RNA modifications using *NanoConsensus*

*NanoConsensus* reports putative modified regions at per-transcript level using as input the outputs obtained from each of the 4 different RNA modification detection algorithms previously described (EpiNano, Nanopolish, Tombo and Nanocompore).

In a first step, *NanoConsensus* converts per-position reported values by each software into normalized Z-Scores across each transcript. These scores are then used to select candidate RNA modified positions for each independent software, which must be higher than the provided user-defined threshold (default value = 5) (**Figure S18**).Then, candidate positions for each individual software previously identified are extended into 5-mers. Flexible overlapping is then performed to identify overlapping k-mers across softwares. The regions supported by two or more softwares are saved as putative modified sites. In a second step, *NanoConsensus* re-scales Z-score values between 0 and 1 across each software’s set of values, to make the score comparable across transcripts (otherwise Z-scores are affected by the coverage of each transcript). Rescaled Z-scores are then converted to a *NanoConsensus score*, which is equal to the median of the rescaled Z-scores across all softwares (NanoConsensus score = median (Z-Score Epinano, Z-score Nanopolish, Z-Score Tombo, Z-Score Nanocompore). Thus, every position across the transcript has an assigned NanoConsensus score. Finally, *NanoConsensus* identifies the differentially modified sites as those having a *Nanoconsensus score* greater than 5 (default settings) times the median per-transcript *NanoConsensus* score. Thus, the threshold varies depending on the background signal of the transcript, i.e., if the data is “noisier” (often the case when coverage is low) the *Nanoconsensus* score threshold to identify a site as “differentially modified” within that given transcript will be higher. We should note that this threshold can be modified by the user. All benchmarking results included in this work used threshold=5 (default) throughout all datasets and RNA modification types. To capture modest variations in stoichiometry, the user might prefer to decrease the threshold to 3.5-4.

*NanoConsensus* will produce the following files as outputs: i) a flat file with all raw results obtained by all softwares, ii) a flat file with the putative modified sites according to *NanoConsensus;* iii) BED tracks that can be loaded into IGV to visualize the results in a user-friendly manner; and iv) a PDF file showing both the performance of each algorithm, represented in the form of Z-scores, and *NanoConsensus* scores along the analyzed transcripts. All code has been integrated into the *mop_consensus* module (RNA modification detection module) that is part of the *MasterOfPores* version 2 (MoP2) Nextflow workflow (https://github.com/biocorecrg/MoP2) ^80^.

### Benchmarking RNA modification predictions across datasets with known stoichiometry and coverage

For each modification type included in the benchmarking, reads from wild type *E. coli* were assumed to be 100% modified, and the respective knockout rRNA reads were assumed to be 0% modified for that position (**Table S3**), respectively. For each dataset analyzed, subsampled datasets were generated using one of the two approaches: i) subsamples contained a fixed amount of reads or; ii) samples contained a fixed number of modified reads (both scripts available at GitHub: https://github.com/novoalab/NanoConsensus/scripts/Downsampling). All benchmarking datasets only included full-length reads, defined as reads whose initial mapping position was 50 or lower and their final position was equal or larger than 1525 and 1779, for 16S and 18S transcripts respectively (available at GitHub: https://github.com/novoalab/NanoConsensus/scripts/Full_Length/Extract_FullLength_IDs.sh).

To compare datasets with different modification stoichiometry levels, we performed two complementary analyses using the two above-mentioned subsampling approaches. In the first subsampling approach, each subsampled dataset comprised the same number of reads (n=1000). Then, to achieve different stoichiometry levels (100, 75, 50, 40, 30, 20 and 10%), a specific proportion of randomly selected modified and unmodified full-length reads were mixed. Triplicates per modification type and stoichiometry were built. In the second subsampling approach, the subsampled dataset were composed of triplicates that have a fixed number of *modified* reads. Then, for each stoichiometry level, an increasing amount of unmodified reads were added sequentially. As a result, the population of modified reads was kept constant whereas the total number of reads varied across the samples from the same replicate. Additionally, four full datasets with a different number of modified reads in the 100% modified sample (50, 100, 500 and 1500 reads) were generated. All samples were compared against 1000 full-length reads from their respective *knock-out* strain using *mop_mod* with default parameters (see previous section).

### Detection of RNA modifications from bacterial samples exposed to antibiotics

Bacterial DRS sequencing runs (**Table S2**) were filtered to keep only full-length reads using in-house scripts (all available at GitHub: https://github.com/novoalab/NanoConsensus/scripts/Full_Length), which were defined as reads that covered the full transcript, leaving up to a maximum of 50nt uncovered in the 5’ and/or 3’ ends. In other words, 16S rRNA reads that were kept covered at least from position 50 to 1525, and reads that aligned to the 23S rRNA covered at least from position 50 to 2894. Replicates per condition and time point were included in the analysis (**Figure S19**). Pairwise comparisons of antibiotic-treated and untreated samples were analysed using the MasterOfPores (MoP2) module *mop_mod*, with default parameters, followed by NanoConsensus (also integrated in MoP2), with the following parameters across all transcripts: *--MZS_thr 3.75 --NC_thr 4*. The directionality of the changes in stoichiometry at the identified ‘differentially modified’ rRNA sites was assessed by comparing the sum of basecalling errors of treated and untreated samples using EpiNano (version 1.1.1). We should note that *Nanoconsensus* can be run comparing treated samples to untreated (t=0h) **(Figure S20A)**, but the signal-to-noise ratio improves if matched untreated time points are used (t=1h or t=16h, respectively) (**Figure S20B**).

### Clustering of full length reads based on basecalling errors

To identify differentially modified populations of RNA molecules upon antibiotic exposure, per-read basecalling errors (mismatch, insertion and deletion frequency) were retrieved using EpiNano (v 1.0) ^49^, which reports per-read base-calling errors. For each read, basecalling errors of the 5-mer region centered in each ‘differentially modified’ site identified by NanoConsensus were extracted, and used for downstream analyses. For each read, the total summed error per-read was calculated. The per-read sum of ‘errors’ provides an approximate measure of the modification levels of a given read in those selected regions, relative to the rest of reads. Reads were binned based on their total summed score, and the fraction of reads belonging to each bin, for each treated sample (str-treated, ksg-treated, untreated) was calculated.

### Growth curves of bacterial strains lacking individual methyltransferases

*E. coli* starter cultures of multiple strains (BW25113, *rsmA* KO, *rsmG* KO, *rsmD* KO, *rsmB* KO, *rsmJ* KO, *rsmH* KO and *rsmE* KO) were grown overnight in 10mL tubes at 37°C 180 rpm, in biological duplicates. Overnight cultures were added into a final volume of 200μl in 96-well plates with an initial OD_600_ of 0.01. Different antibiotic concentrations (streptomycin: 8, 16, 32, 64 and 128μl/mL; kasugamycin: 62, 125, 250, 500 and 1000μl/mL) were tested for each strain. For each strain and antibiotic concentration, three technical replicates were generated and the experiment was performed with two biological replicates, in two independent days. Bacteria were grown at 37°C and OD_600_ was measured every 10 minutes with a TECAN M200 Plate Reader. Area Under the Curve (AUC) calculations were performed in R using the *growthcurver* package ^90^.

### Bacterial ribodepletion

Ribodepletion was performed using riboPOOL oligos (siTOOLs, cat #055) following the manufacturer’s protocol (version: riboPOOL Protocol_v1-8). Briefly, 5ug of total RNA was mixed together with 1uL resuspended riboPOOL oligos, 5uL hybridization buffer (10 mM Tris-HCl pH 7.5, 1 mM EDTA, 2 M NaCl) and 0.5 uL SUPERase•In RNase Inhibitor (Thermo Fisher, AM2694) followed by a 10 minute incubation at 68°C and slow cool down to 37°C for hybridization. In parallel, Dynabeads MyOne Streptavidin C1 (Thermo Fisher, #65001) beads were resuspended, and 80uL of the resuspended beads were transferred into a tube and placed on a magnetic rack. Then, the supernatant was removed and 100 uL of bead resuspension buffer (0.1 M NaOH, 0.05 M NaCl) was added to the sample. Beads were then resuspended in the buffer by agitation, the tube was placed again on a magnetic rack, and the supernatant was aspirated. This step was performed twice. Beads were then resuspended in 100uL of washing buffer (0.1 M NaCl) and placed again onto the magnet to remove the supernatant. The beads were then resuspended in 160uL of depletion buffer (10 mM Tris-HCl pH 7.5, 1 mM EDTA, 1 M NaCl). This mixture was then divided into two tubes of 80uL, which were used sequentially. 20uL of hybridized riboPOOL oligos and total RNA was spun down briefly and then transferred into a tube with 80uL of beads in depletion buffer. Then, it was mixed by pipetting carefully and incubated at 37°C for 15 minutes, followed by a 50°C incubation for 5 minutes. The reaction was placed on the magnetic rack and the supernatant was transferred into the second tube of beads, from which the supernatant was removed before its use. The solution was mixed and incubated at 37°C for 15 minutes, followed by a 50°C incubation for 5 minutes. After briefly spinning down the droplets, the mix was placed on a magnet for 2 minutes and the supernatant was transferred into a different tube. Finally, long RNA species (>200bp) were separated from short ones (<200) using RNA Clean & Concentrator-5 kit (Zymo, R1013).

### End-capture nanopore cDNA sequencing (Nano3P-seq)

*E. coli* total RNA was sequenced using Nano3P-seq ^79^, which is a cDNA library preparation variation that relies on the SQK-DCS109 direct cDNA nanopore sequencing kit. Notably, Nano3P-seq does not require the presence of polyA tails, which are typically absent in bacterial mRNAs, to initiate the library (whereas the standard direct cDNA nanopore sequencing library requires the presence of polyA tail). The protocol was executed as previously described (https://www.protocols.io/view/nano3p-seq-protocol-3byl4j292lo5/v1). Briefly, 50ng of ribodepleted total RNA per sample was mixed with 1 µL pre-annealed RNA-DNA oligos, 1uL 100 mM DTT, 4 µL 5X TGIRT buffer, 1uL RNasin Ribonuclease Inhibitor (Promega, N2511), 1uL TGIRT (InGex) and nuclease-free water (NFW) up to 19uL.This reverse transcription mix was first incubated at RT for 30 minutes before 1uL 10 mM dNTP mix was added. Then, the solution was brought to 60°C for 60 minutes, and inactivated by heating at 75°C for 15 minutes before moving to ice. Afterwards, RNAse Cocktail (Thermo Scientific, AM2286) was added followed by an incubation at 37°C for 10 minutes and a clean up step using 0.8X AMPure XP Beads (Agencourt, A63881). Then, 1 µL 100 µM of the DNA oligo, complementary to the initial one, was added to 15uL solution containing cDNA with 2.25 µL 0.1 M Tris pH 7.5, 2.25 µL 0.5 M NaCl and 2 µL NFW. The mix was incubated at 94°C for 1 minute and the temperature was ramped down to 25 °C (−0.1°C/s). Later, 2.5uL from a single native barcode and 25 µL Blunt/TA Ligase Mix (NEB, M0367S) were added, followed by a 10 minutes incubation at RT and a clean up step with 0.5X AMPure XP beads and eluted with 16uL NFW. The concentration of each sample was quantified using Qubit ssDNA HS assay (Thermo Scientific, Q10212). Individual samples were pooled together with a final concentration that did not exceed 200 fmol and a final volume of 65uL. All the subsequent steps used reagents from the direct cDNA nanopore sequencing kit (ONT, SQK-DCS109). The pooled sample was mixed with 5uL Adapter Mix (AMI), 20uL 5X NEB Quick Ligation buffer (NEB, B6058S) and 10uL Quick T4 DNA ligase (NEB, M0202M) and then incubated for 10 minutes at RT. Finally, the sample was cleaned with 0.5X AMPure XP beads (Beckman Coulter, A63881), using washing buffer (WSB) and elution buffer (EB). The library was mixed with sequencing buffer (SQB) and loading Beads (LB) prior to loading it onto a primed R9.4.1 MinION flowcell. Biological triplicates were run on independent MinION flowcells, and prepared on different days.

### Base-calling, demultiplexing and mapping of Nano3P-seq datasets

FAST5 reads sequenced with the Nano3Pseq protocol were analyzed using the *MasterOfPores* version 2 (MoP2) Nextflow workflow ^61^ (**Table S3**). Firstly, the *mop_preprocess* module was used to basecall and demultiplex all FAST5 using Guppy 4.0 (Oxford Nanopore Technologies) with the DNA basecalling model *dna_r9.4.1_70bps_hac*. Reads whose barcode could not be identified, went through a second round of demultiplexing using readucks (version 0.0.3) ^91^ with parameters *--limit_barcodes_to 1 2 3 4 5 6 --adapter_threshold 73 --threshold 50*. Demuxed reads were then aligned to the *E.coli BW251113* genome (NCBI ID: CP009273.1) using minimap2 v.2.17 with -ax spliced -k14 -uf parameters. Both per-gene counts and per-transcript counts were generated. Briefly, per-gene counts were obtained by htseq-count (version 0.13.5) ^92^ with the option --stranded reverse as well as using the *--nonunique all* option to account for reads spanning more than one feature (ie: bacterial operons). On the other hand, to obtain per-transcript estimates, bacterial reads were aligned to the reference transcriptome that was built from the reference genome annotation (NCBI ID: CP009273.1, see code in GitHub: https://github.com/novoalab/NanoConsensus/Nano3Pseq_analysis/) using minimap2 v.2.17 with *-ax map-ont -k14* parameters. Per-transcript abundance estimates were obtained from the aligned reads using salmon quant (version 1.9.0) ^93^ with the parameters *--ont -l SR*. Both per-gene counts (using either –stranded and per-transcript estimates were used as input for differential expression analysis (**Figure 5C**, see also **Figure S21**).

### Differential expression and GO term enrichment analysis

Differential expression analysis was performed both on per-gene and per-transcript counts of untreated samples compared to antibiotic-treated samples (streptomycin or kasugamycin) using DeSeq2 v.1.34.0 ^94^. Genes with an absolute log2(Fold Change) greater or equal than 2 and an adjusted p-value lower or equal than 0.05 were considered as differentially expressed. Results were visualized using the R package *EnhancedVolcano* v.1.12.0 ^95^ (**Figure 5C** and **Figure S21**). Scripts used for differential expression analysis are available in GitHub: https://github.com/novoalab/NanoConsensus/scripts/Differential_Expression). Genes with absolute log2(Fold Change) greater or equal than 2 were used as input for a GO term enrichment analysis, which was performed using Panther (release 17.0) ^96^. All genes included in the differential expression analysis were included as the reference list for this analysis. The annotation data set used was GO biological process complete, the statistical test was the Fisher’s Exact Test, followed by calculating the False Discovery Rate (FDR). Only terms with a FDR<0.05 were included in the final results (**Figure 5D**).

## CODE AVAILABILITY

*NanoConsensus* code and documentation is publicly available in GitHub (https://github.com/novoalab/NanoConsensus) and in Zenodo (https://doi.org/10.5281/zenodo.5805806) Moreover, to facilitate its implementation by future users, *NanoConsensus* has been integrated as a module into the *MasterOfPores* ^61^ Nextflow ^97^ workflow for the analysis of direct RNA nanopore sequencing data (https://github.com/biocorecrg/MoP2), under version 2.0 of *MasterOfPores*.

## DATA AVAILABILITY

Base-called FAST5 files have been deposited in the European National Archive under the accession codes PRJEB42568 (*E. coli* total RNA DRS samples from different strains), PRJEB48806 (*E. coli* total RNA DRS samples exposed to antibiotics) and PRJEB58640 (*E. coli* Nano3P-seq ribodepleted total RNA samples). Base-called FAST5 files from yeast total RNA were taken from previously published data, available at ENA under accession PRJEB37798 ^72^. A description of all samples used in this work with their corresponding ENA numbers can be found in **Tables S2-S4**.

## AUTHOR CONTRIBUTIONS

AD-T performed most wet lab experiments and data analyses described in this work, including the development, implementation and benchmarking of NanoConsensus. RM helped with the *E. coli* growth curve experiments. OB helped with preparing and sequencing some of the direct RNA libraries used in this work. LC implemented NanoConsensus into a Nextflow module and into the *MasterOfPores* general workflow. JP supervised the implementation of NanoConsensus into *MasterOfPores*. XB built structural superimpositions of ribosome structures used for 3D analysis of rRNA modifications relative to antibiotic and tRNA binding sites. EMN conceived and supervised the work. AD-T built the figures. AD-T and EMN wrote the manuscript, with contributions from all authors.

## ACKNOWLEDGEMENTS

We thank all the members of the Novoa lab for their insightful discussions. We thank Prof. Xavier Barril for his discussions and help with Pymol to obtain the structural alignments of PDB structures of ribosomes from different species containing diverse antibiotics. AD-T is supported by an FPI Severo-Ochoa fellowship by the Spanish Ministry of Economy, Industry and Competitiveness (MEIC). This work was supported by funds from the Spanish Ministry of Economy, Industry and Competitiveness (MEIC) (PID2021-128193NB-100 to EMN) and the European Research Council (ERC-StG-2021 No 101042103 to EMN). We acknowledge the support of the MEIC to the EMBL partnership, Centro de Excelencia Severo Ochoa and CERCA Programme / Generalitat de Catalunya.

## COMPETING INTERESTS

EMN is a member of the Scientific Advisory Board of IMMAGINA Biotech. EMN has received travel and accommodation expenses to speak at Oxford Nanopore Technologies conferences. AD-T and OB have received travel bursaries from ONT to present their work in conferences. Otherwise, the authors declare that the research was conducted in the absence of any commercial or financial relationships that could be construed as a conflict of interest.

