## Supplementary Information - Figures S1-S21 for "Native RNA nanopore sequencing reveals antibiotic-induced loss of rRNA modifications in the A- and P-sites"

### SUPPLEMENTARY FIGURES

**Figure S1. Lack of m5C can be detected in the form with direct RNA nanopore sequencing. (A)** IGV snapshots of *E.coli* WT and knockout strains illustrating loss of base-calling errors upon *rsmF* knockout. $m^5C$ modifications cause increased deletions and insertions at position +1, in agreement with previous works [1], and loss of m5C (*rsmF* knockout) shows decreased deletion frequency at position +1. Positions with mismatch frequencies greater than 0.1 are coloured, whereas positions with mismatch frequencies lower than 0.1 are shown in gray. **(B)** Scatterplot of the summed base-calling error frequencies (sum of insertion, deletion and mismatch frequencies) at each nucleotide position in the knockout strain, relative to WT. The rRNA modified sites that are lost upon knockout of the gene are shown in red (position 0); the neighboring positions (±4 nt) to the rRNA modification site (position 0) are shown in blue. Remaining positions are shown in gray.

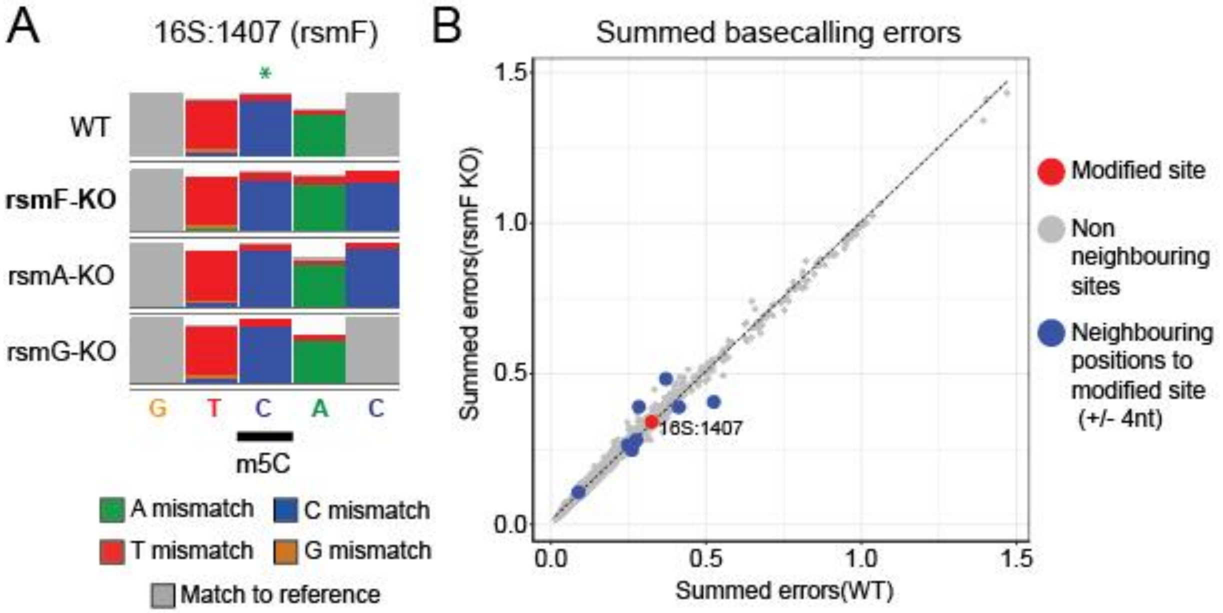

**Figure S2. Raw score tracks depicting differential RNA modification levels predicted by each individual software when comparing untreated and antibiotic-treated samples. (A,B)** Per-position raw scores (ΔSummed errors for EpiNano; ΔMedian current intensity for Nanopolish; -log10(p-value) for both Tombo and Nanocompore) assigned by each software when comparing untreated and 1h streptomycin-treated (A) or kasugamycin-treated samples (B). In grey, non-significant positions; and, in color, positions reported as modified by every algorithm which are the same as the ones described in [2]**.** Data from two independent biological replicates is shown.

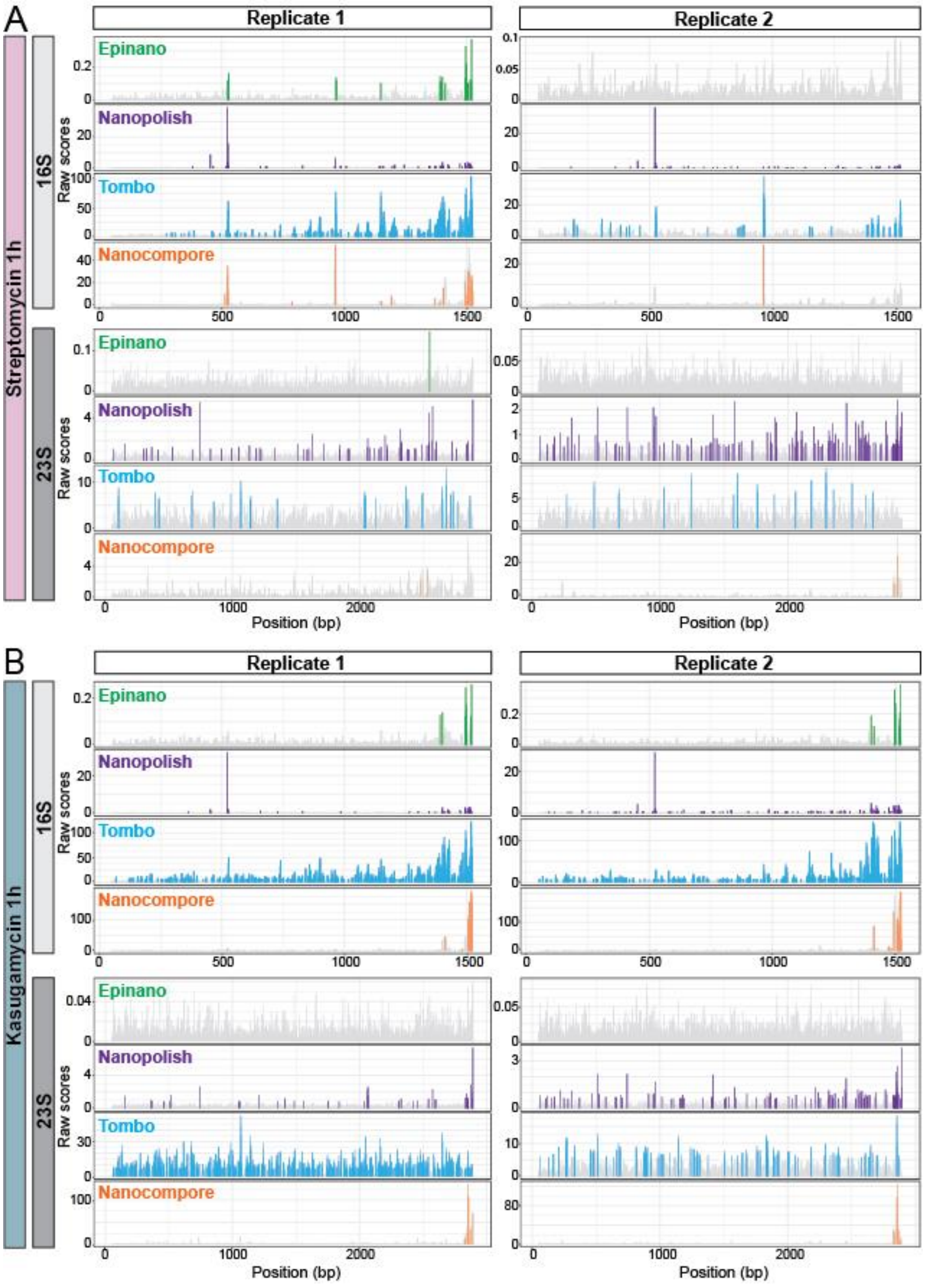

**Figure S3. Z-Score tracks depicting differential RNA modification levels predicted by each individual software when comparing untreated and streptomycin-treated samples. (A,B)** Z-score values of scores obtained by each softwares along 16S and 23S after exposure to streptomycin for 1h (A) or 16h (B) at a concentration of 16µl/mL. Data from two independent biological replicates is shown.

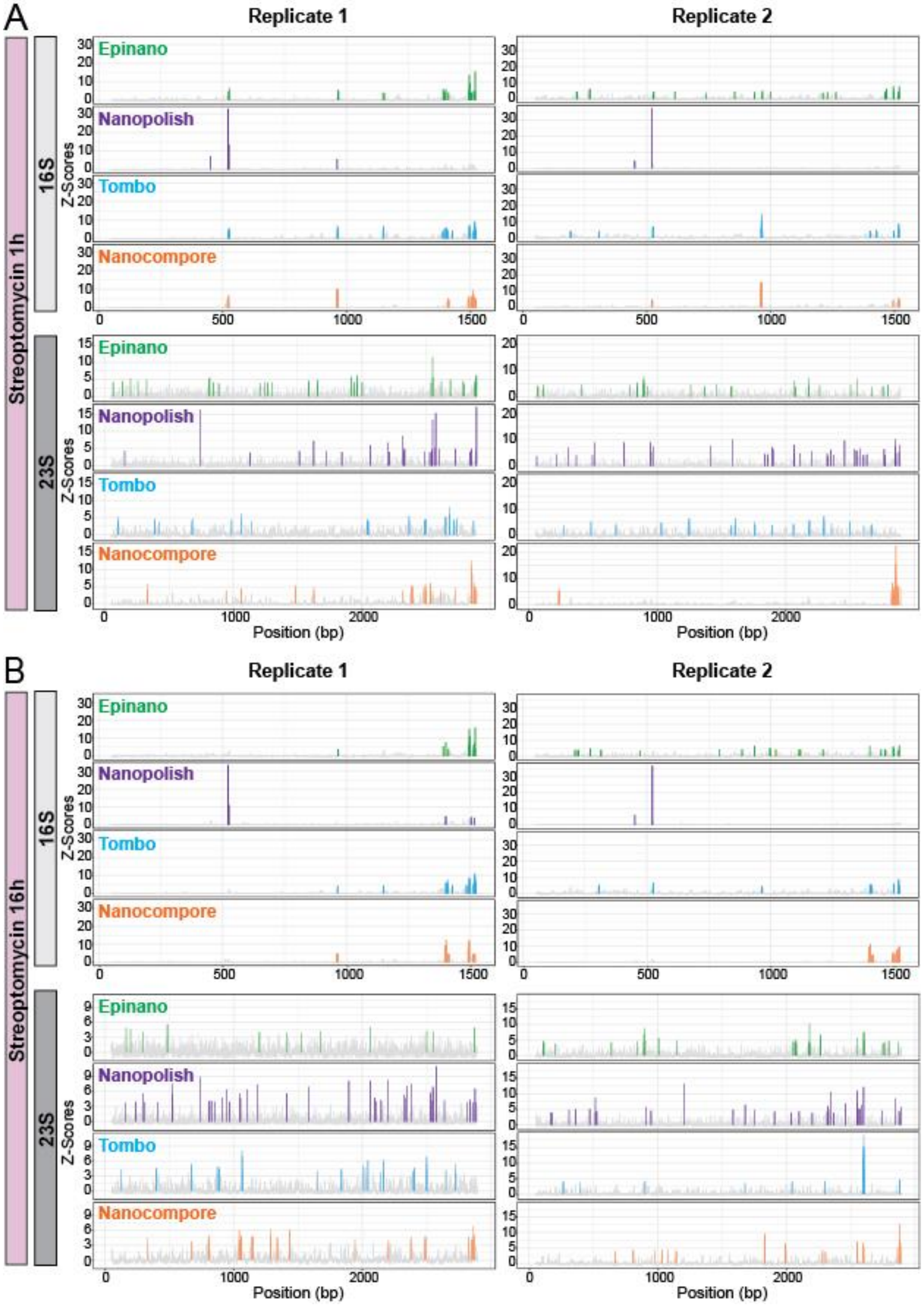

**Figure S4. Z-Score tracks depicting differential RNA modification levels predicted by each individual software when comparing untreated and kasugamycin-treated samples. (A,B)** Z-score values of scores obtained by each softwares along 16S and 23S after exposure to kasugamycin for 1h (A) or 16h (B) at a concentration of 250µl/mL. Data from two independent biological replicates is shown.

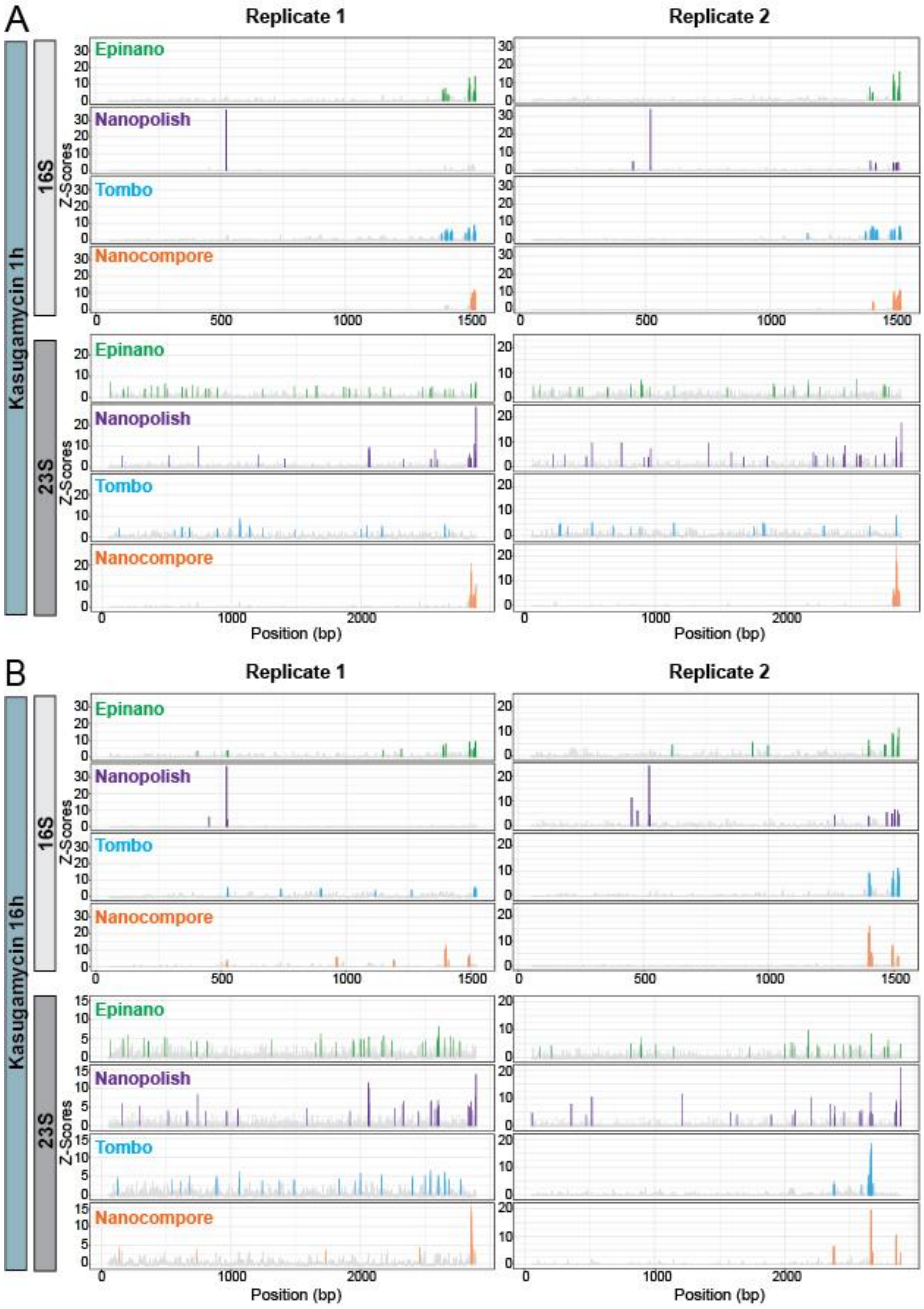

**Figure S5. Overlap of predicted differential RNA modified sites across softwares. (A)** Venn Diagrams showing the predicted differentially modified sites by each software across replicates. Data from both antibiotic exposures (streptomycin and kasugamycin), time points (t=1h and t=16h) and transcripts is included in the plots. Predicted sites in replicate 1 are shown in full lines, whereas predicted sites in replicate 2 are shown in dashed lines. **(B)** Venn Diagrams showing the replicability of predicted sites across softwares. Data from both antibiotic exposures (streptomycin and kasugamycin), time points (t=1h and t=16h) and transcripts is included. Intersection % refers to the sites predicted in both replicates divided by the total number of sites.

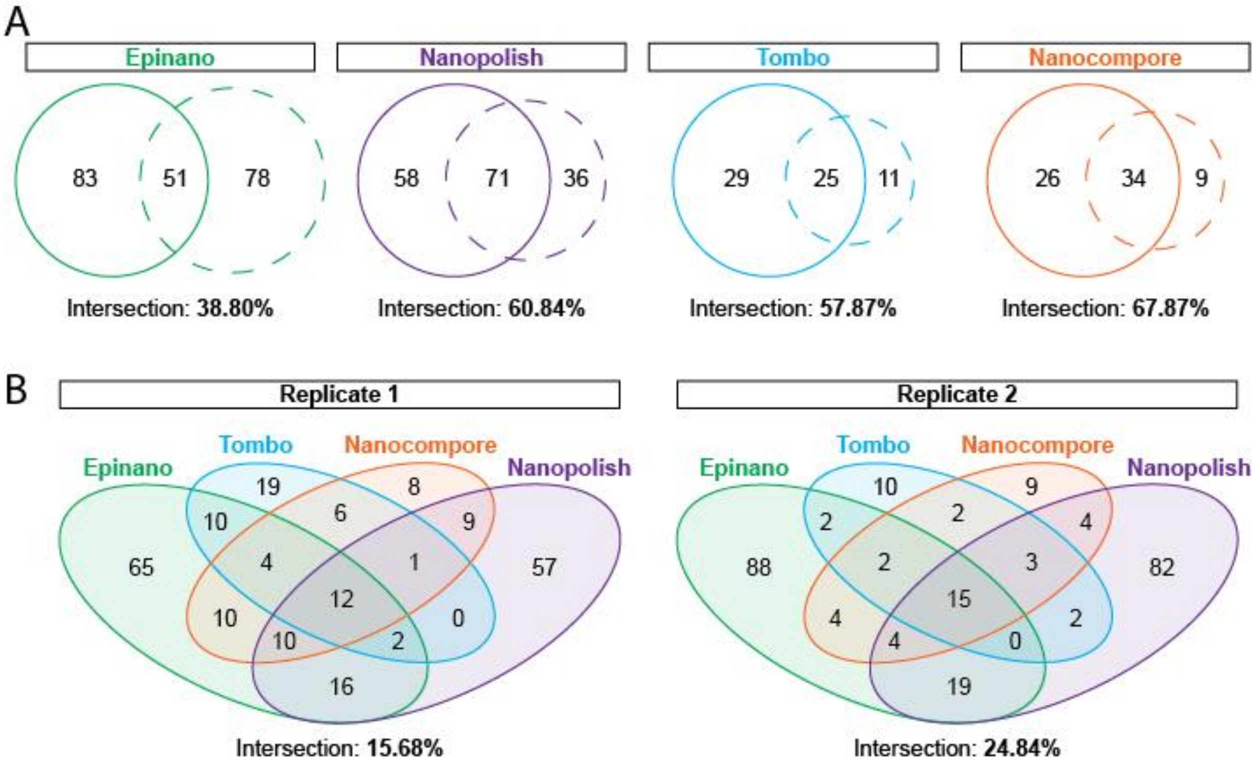

**Figure S6. Softwares' performance when comparing 50 reads samples. (A)** Z-values of scores given by all softwares across the *E. coli* 16S rRNA when using 50 reads as input for each condition. Modified sites are indicated with an asterisk (*). Data from three independent replicates is shown. **(B)** Venn Diagrams showing the replicability of predicted sites across softwares. Data from all comparisons is included.

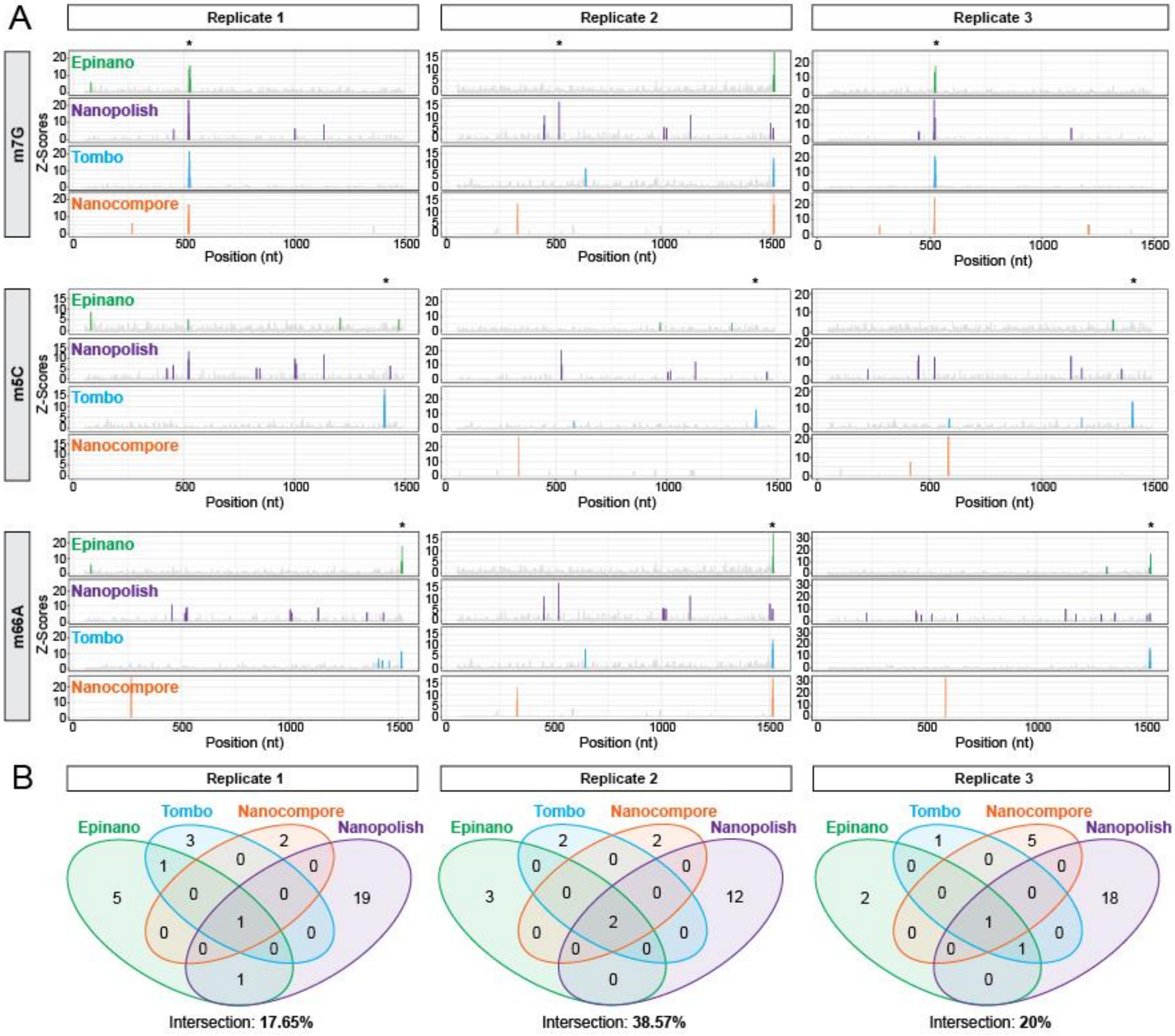

**Figure S7. Softwares' performance when comparing 100 reads samples. (A)** Z-values of scores given by all softwares across the *E. coli* 16S rRNA when using 100 reads as input for each condition. Modified sites are indicated with an asterisk (*). Data from three independent replicates is shown. **(B)** Venn Diagrams showing the replicability of predicted sites across softwares. Data from all comparisons is included.

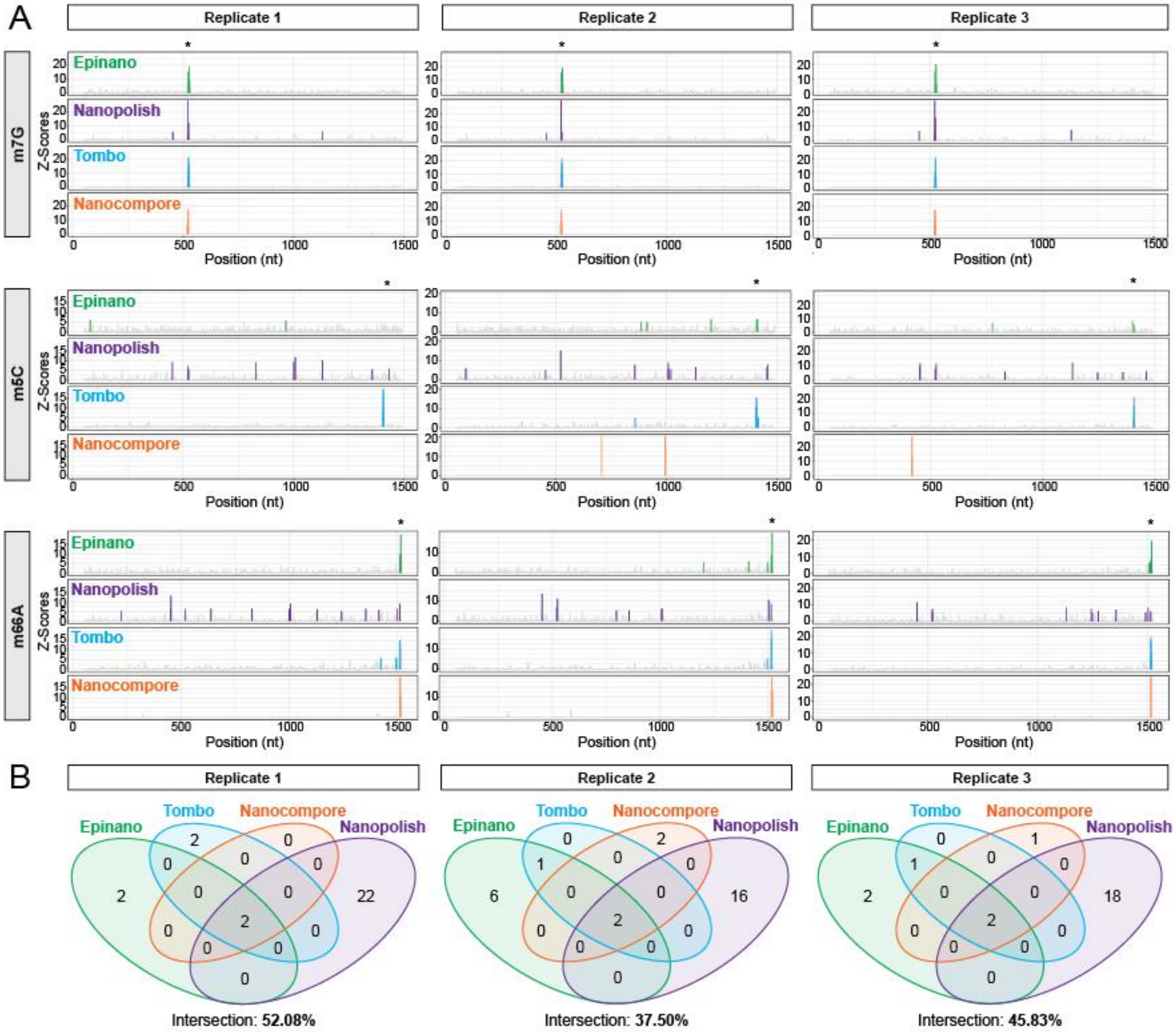

**Figure S8. Softwares' performance when comparing 500 reads samples. (A)** Z-values of scores given by all softwares across the *E. coli* 16S rRNA when using 500 reads as input for each condition. Modified sites are indicated with an asterisk (*). Data from three independent replicates is shown. **(B)** Venn Diagrams showing the replicability of predicted sites across softwares. Data from all comparisons is included.

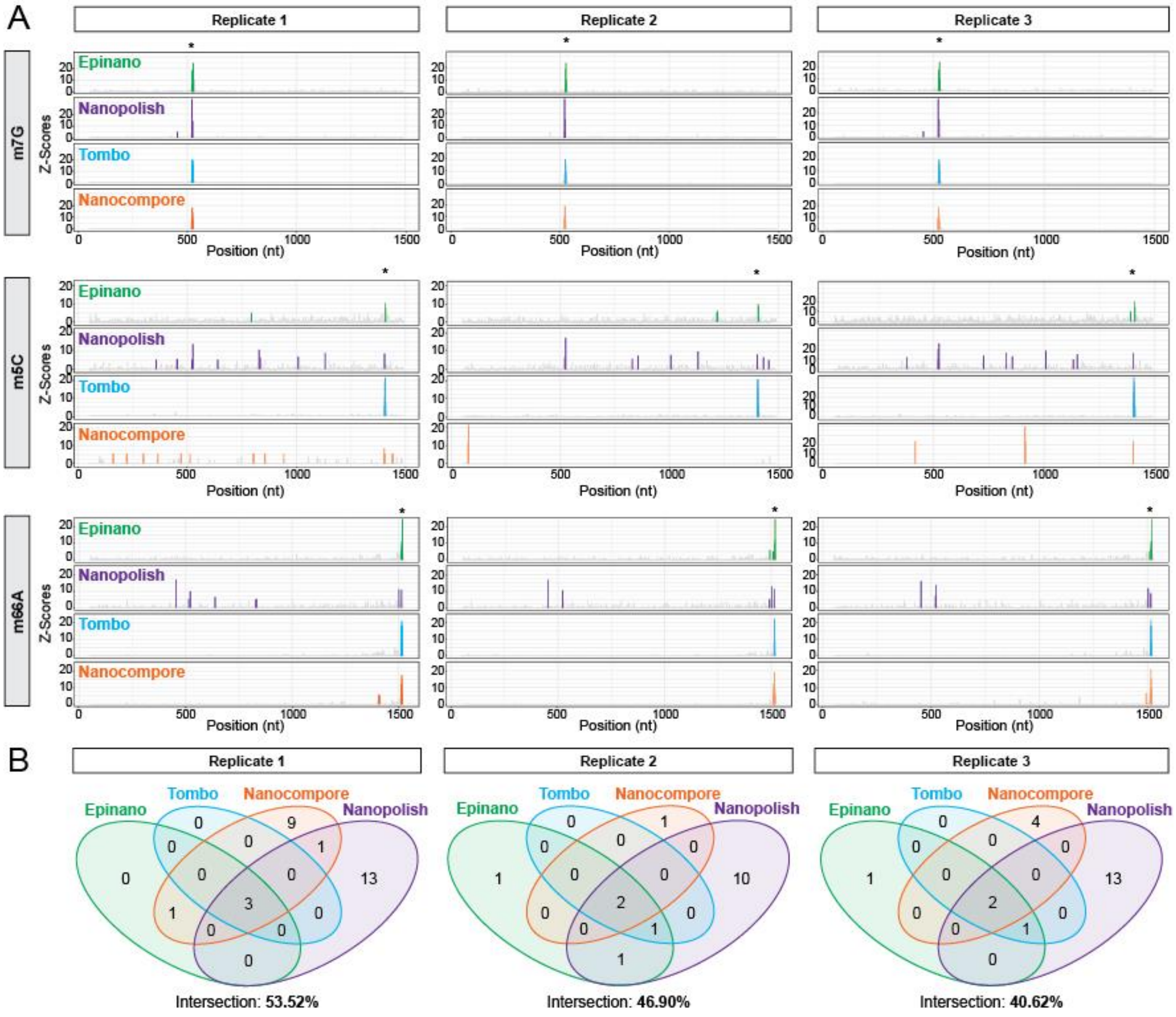

**Figure S9. Softwares' performance when comparing 1500 reads samples. (A)** Z-values of scores given by all softwares across the E. coli 16S rRNA when using 1500 reads as input for each condition. Modified sites are indicated with an asterisk (*). Data from three independent replicates is shown. **(B)** Venn Diagrams showing the replicability of predicted sites across softwares. Data from all comparisons is included.

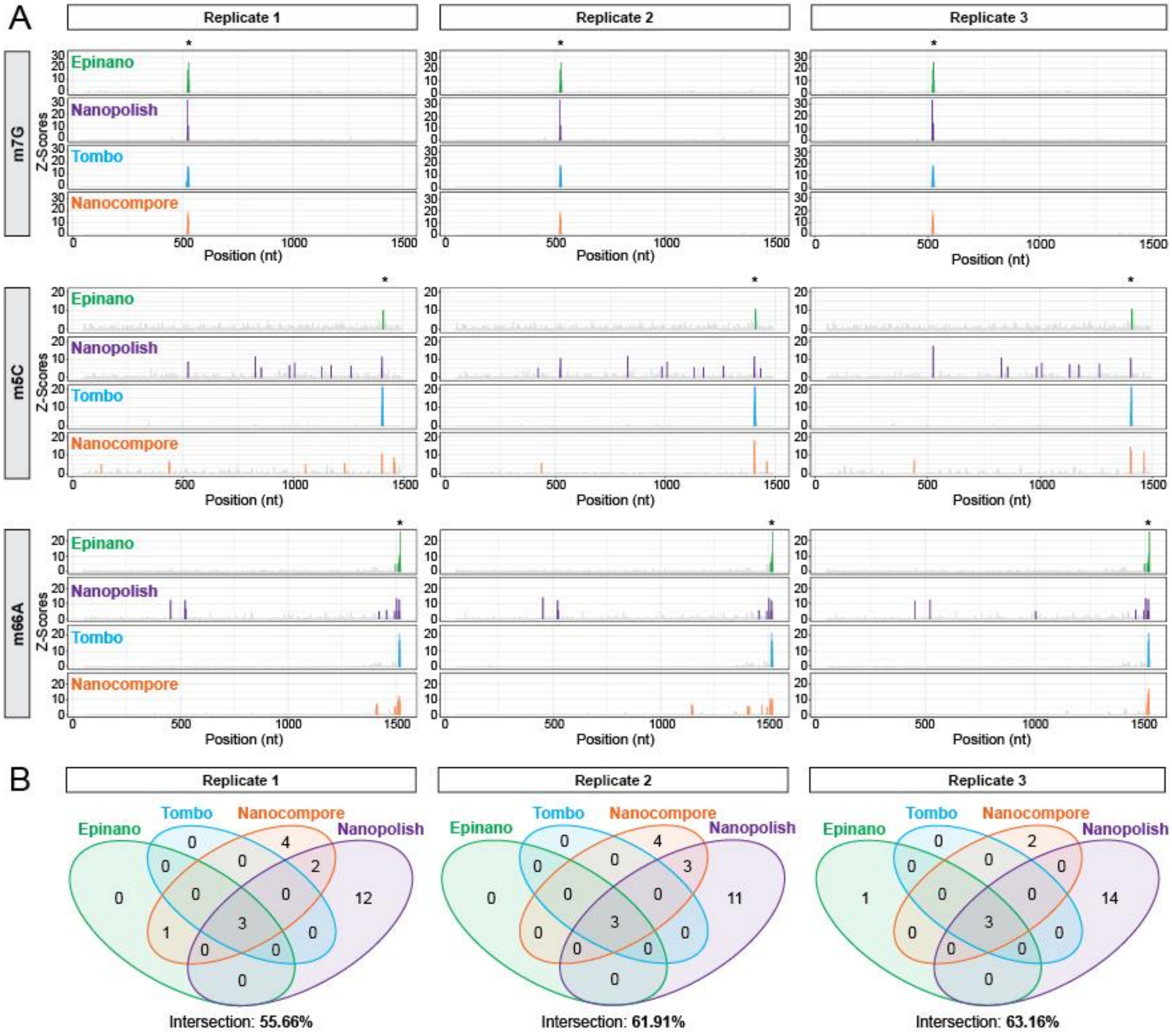

**Figure S10. AUC values and additional ROC curves from the benchmarking procedure. (A)** Boxplots representations of AUC values obtained by each individual software, for each of the 6 modifications studied. All results from all stoichiometry (10, 20, 30, 40, 50, 75 and 100) and coverage levels (50, 100, 500, 1500) have been combined. Box, first to last quartiles; whiskers, 1.5x interquartile range; center line, median; points, outliers. See Table S5 for median AUC values for all conditions included in the study. **(B)** ROC curves from *Nanopolish* (left panels) *Tombo* (middle panels) and *Nanocompore* (right panels) across a range of modification types and coverage levels are shown. Each curve was built using merged data from three independent replicates. At the bottom corner, their respective AUC values are shown. See also Table S5 for median AUC values corresponding to all conditions included in the study.

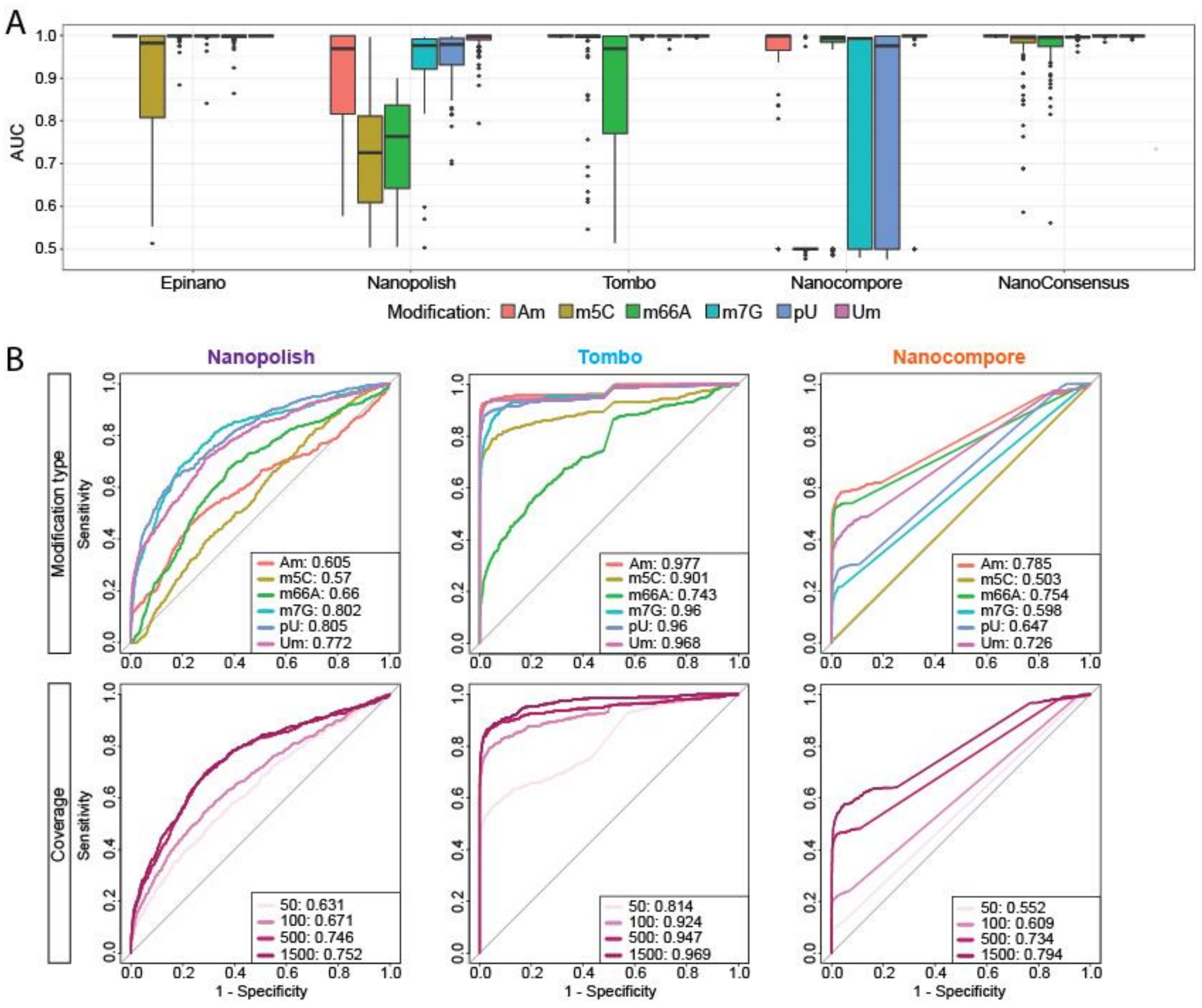

**Figure S11. Additional analysis to assess the performance of each software. (A)** Dotplot of NanoConsensus scores obtained across the modified 5-mer for every modification type present in the benchmarking. Results from all coverage levels in 100% modified samples are included. Box, first to last quartiles; whiskers, 1.5× interquartile range; center line, median. **(B)** Boxplot representations of AUC values obtained by *NanoConsensus* for each of the 6 modifications studied. All results from all stoichiometry (10, 20, 30, 40, 50, 75 and 100). Box, first to last quartiles; whiskers, 1.5x interquartile range; center line, median; points, outliers.

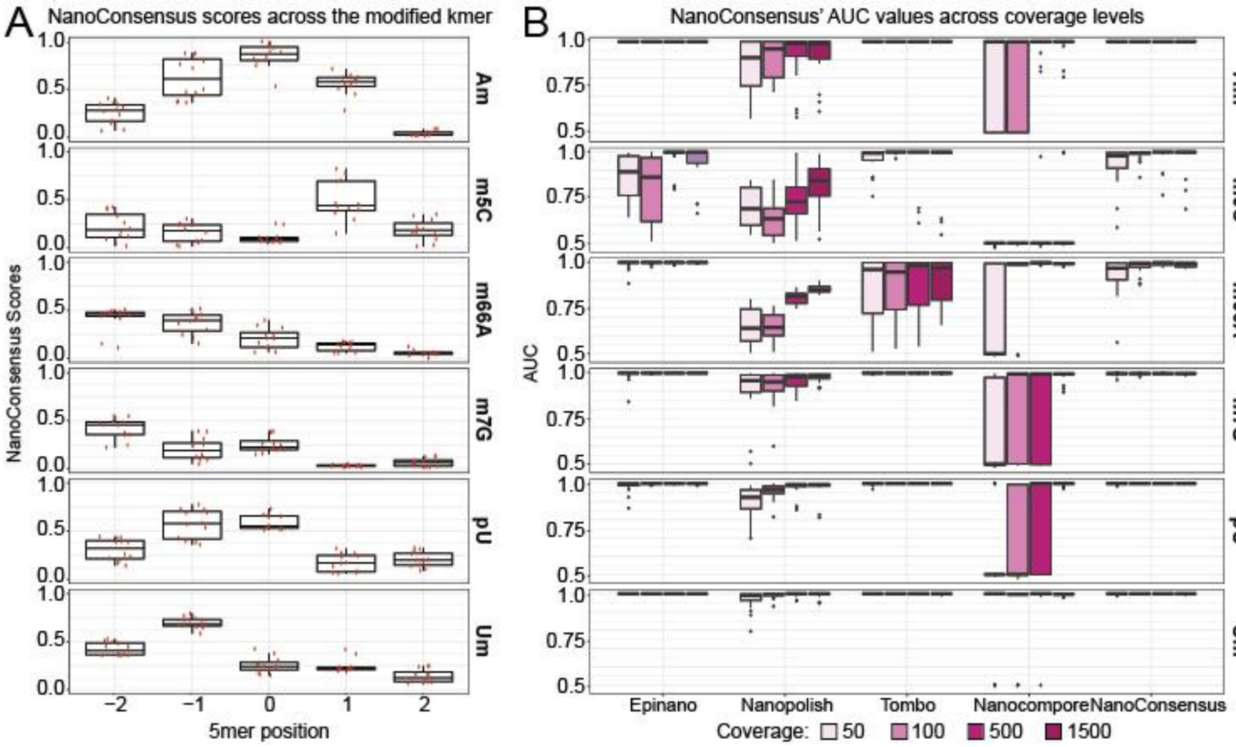

**Figure S12. Radar plots from positive predictive values (PPV) (A)** Radar plots of positive predictive values (PPV) when using a Z-Score threshold > 5 for each software. Values for each coverage, modification, and stoichiometry level are shown. **(B)** Radar plots of positive predictive values (PPV) when using a Z-Score threshold > 5 and a *NanoConsensus* score threshold > 5*(median across transcript) *(see Methods*). Values for each coverage, modification, and stoichiometry level are shown.

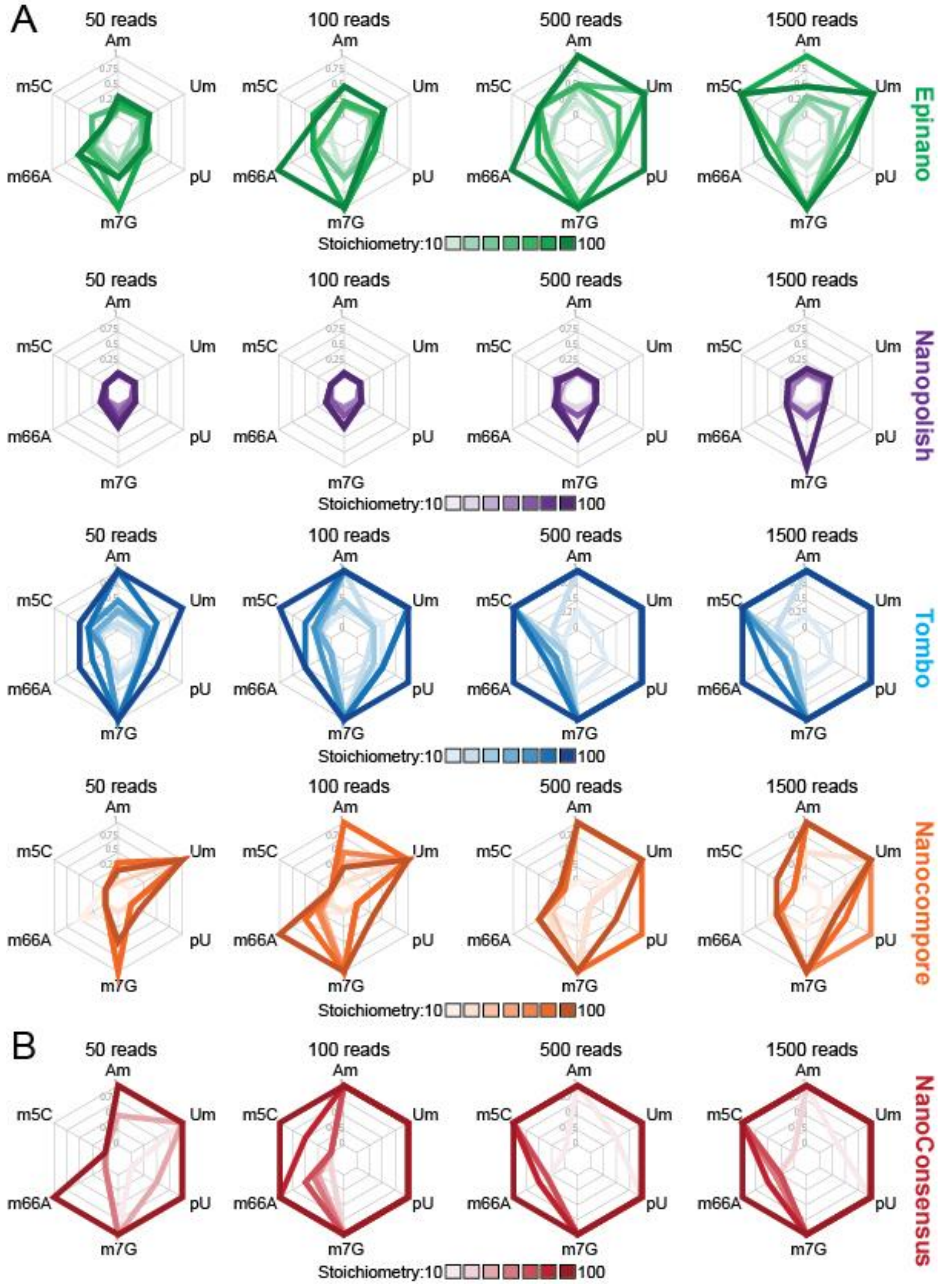

**Figure S13. Decrease of the summed basecalling errors at the modified sites upon antibiotic exposure.** Scatter plots of median basecalled errors from treated and untreated DRS samples across 16S. Summed base-calling errors were obtained using *EpiNano*, which was run as part of the *Nanoconsensus* workflow. Differentially modified sites upon antibiotic exposure identified by *NanoConsensus* are shown in red.

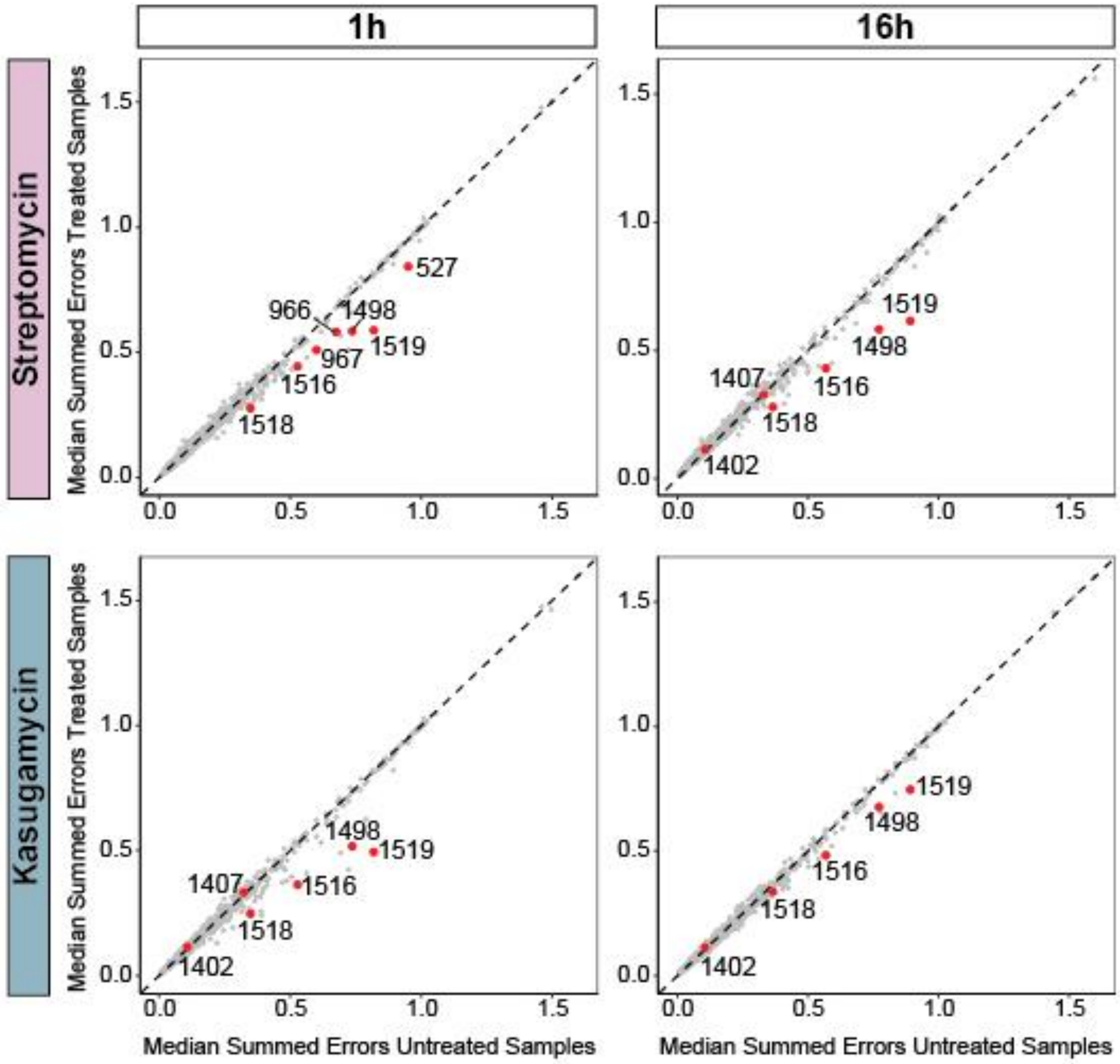

**Figure S14. Principal Component Analysis (PCA) of per-read basecalling errors. (A)** Biplot of scores of PC1 and PC2, depicting per-read base-calling errors, from untreated, str-treated and ksg-treated samples, 1h after antibiotic exposure. Only reads with summed error (SE) lower than 4 in differentially modified regions identified by *Nanoconsensus* are included. Reads are colored by their summed error (proxy for RNA modification levels) in differentially modified kmers upon antibiotic exposure**.** Each dot represents a read. **(B)** PCA loadings from PC1 and PC2, depicting the RNA modified sites that are contributing to the separation of the reads based on their summed errors (proxy for RNA modification levels)

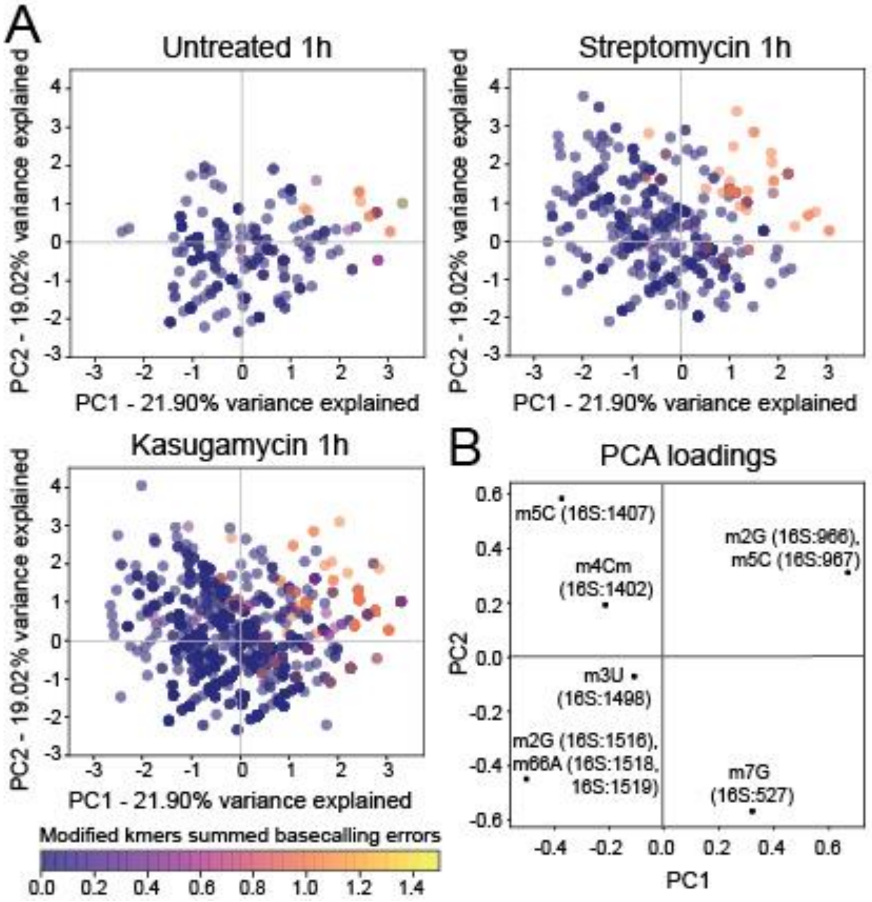

**Figure S15. Growth curves from remaining KO strains.** Median growth curves from four biological replicates of different E.coli knock-out strains (rsmB, rsmD, rsmH and rsmJ) across a range of antibiotic (streptomycin, upper panels; kasugamycin, lower panels) concentrations. Growth curve data from all three technical replicates from each biological replicate per strain and condition has been fitted to a logistic curve.

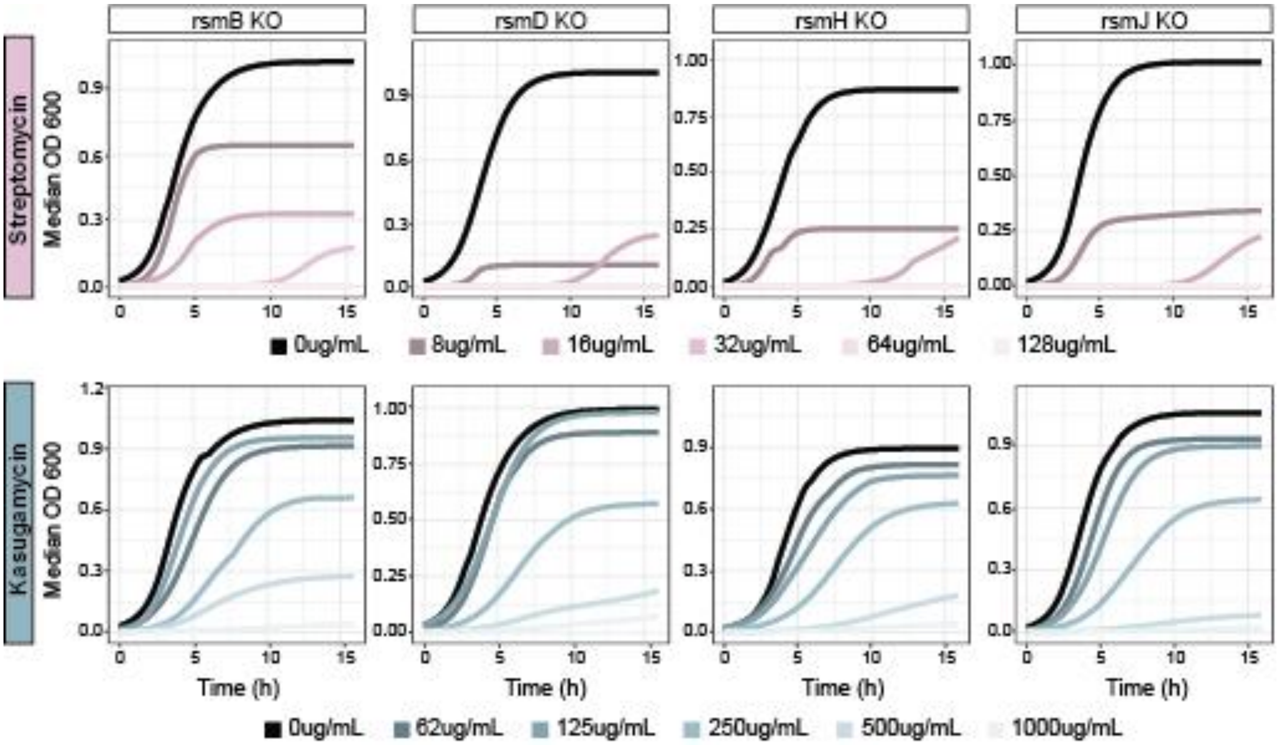

**Figure S16. TapeStation profiles from samples used in this study. (A)** Profiles from total RNA extracted with Trizol coupled with RNeasy Mini columns or standalone (see *Methods*). **(B)** Total RNA profiles performed as quality control during the library preparation. **(C)** Total RNA profiles of in vitro polyadenylated total RNA used as input for direct RNA nanopore sequencing libraries from 4 *E. coli* strains used in this study (BW25113 wild type, *rsmA* knockout, *rmsG* knockout and *rsmF* knockout).

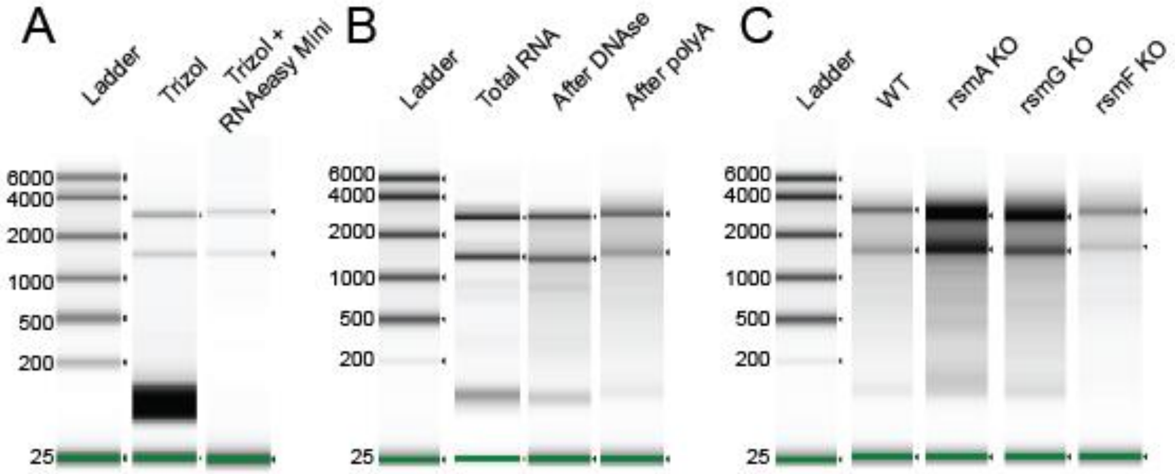

**Figure S17. Overlap of reported modified sites by each algorithm using the thresholds described in** [2] **when comparing wild-type and knock-out bacterial samples. (A)** Venn Diagrams showing the replicability of predicted sites across softwares. **(B)** Venn Diagrams showing the replicability of predicted modified kmers across softwares.

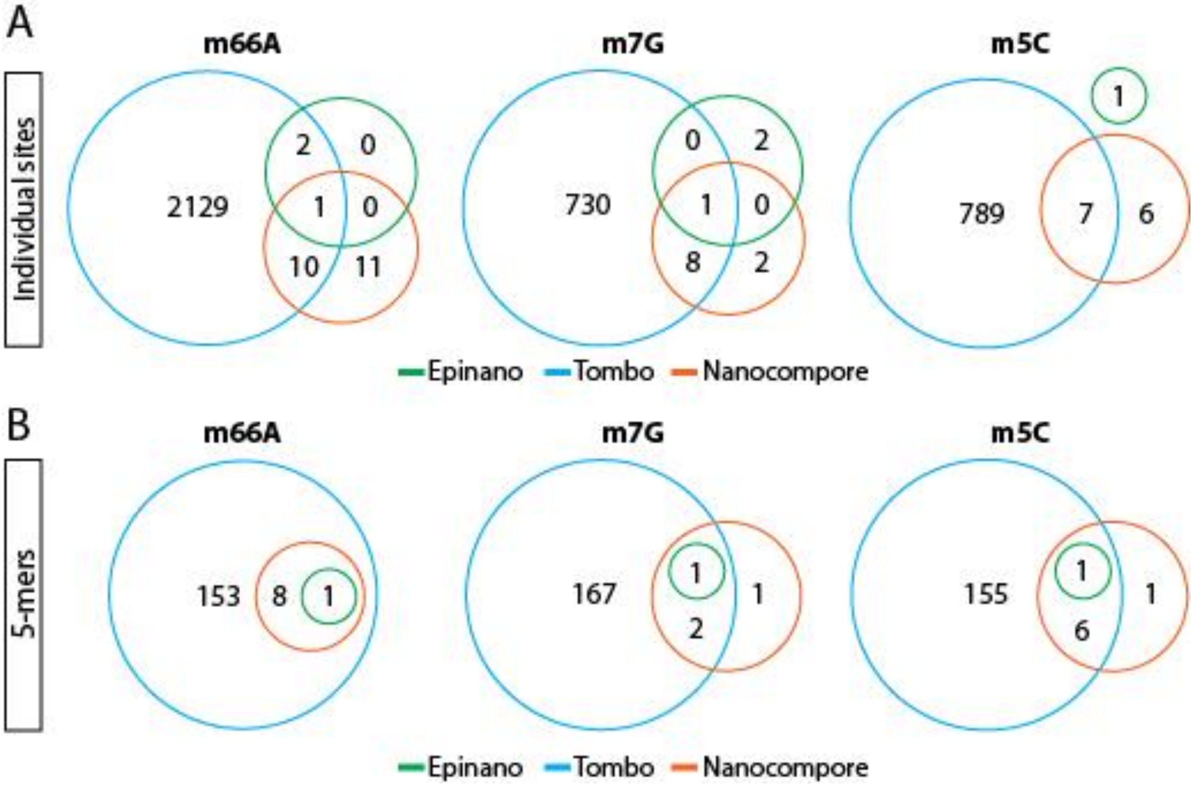

**Figure S18. Z-score score tracks depicting differential RNA modification levels predicted by each individual software when comparing wild-type and knock-out bacterial samples. (A,B,C)** Z-values of scores given by all softwares across the 16S transcript when comparing the WT against each KO strain using all reads from all samples. Modified sites placed by the modification enzyme that is deleted – $m^7$G527 in *rsmG* KO (**A**); $m^5$C1407 in *rsmF* KO **(B)**, $m^{6,6}$A1518 and $m^{6,6}$A1519 in *rsmA* knockout **(C)** – are highlighted with an asterisk (*).

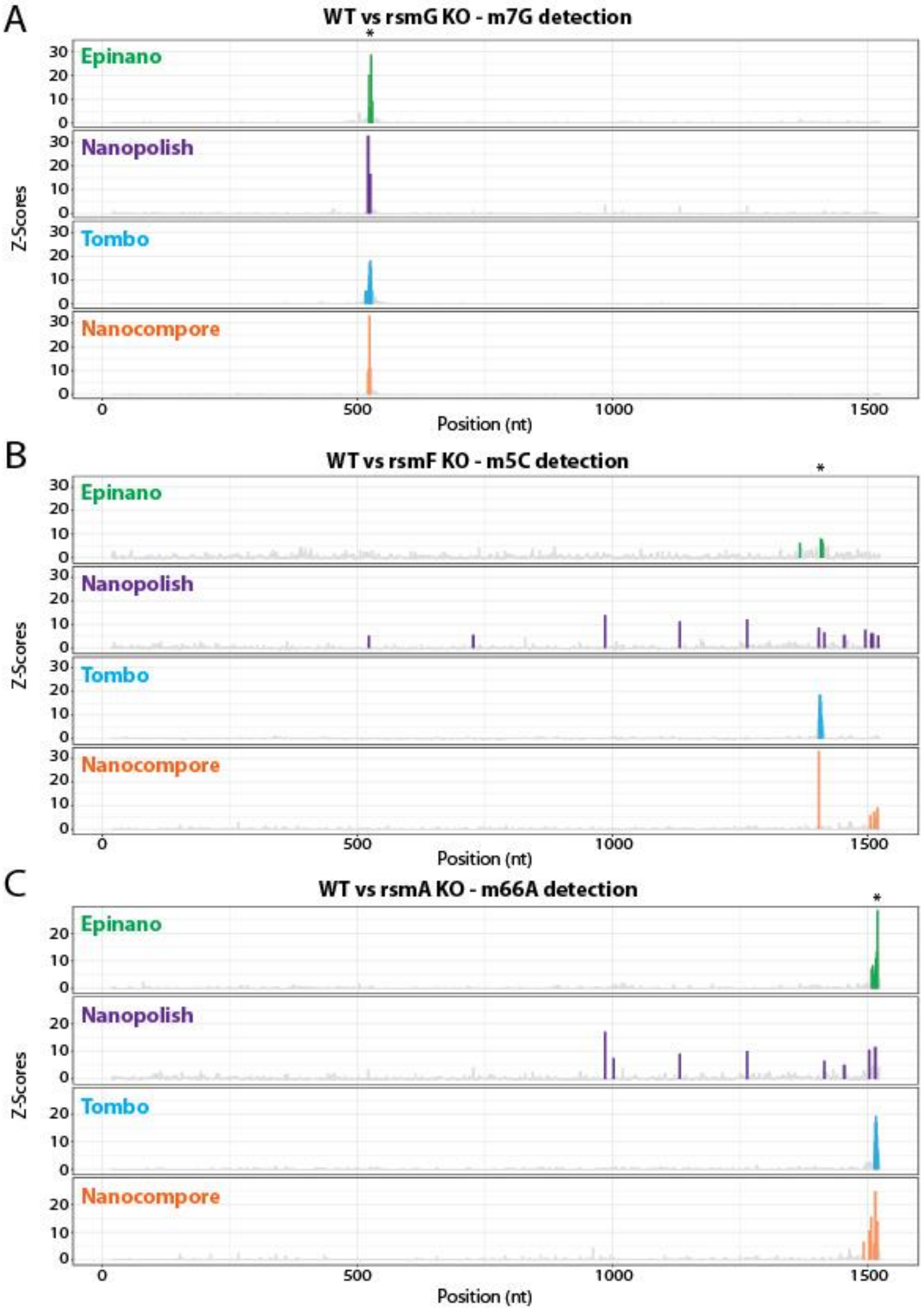

**Figure S19: Replicability of summed errors across biological replicates from both wild-type samples and antibiotic exposed samples.** Scatterplots depicting the replicability of summed base-calling errors (mismatch, deletion and insertion frequency) from streptomycin-treated (left panels) kasugamycin-treated (middle panels) and untreated (right panels) samples, which were collected 1h or 16h post-antibiotic treatment.

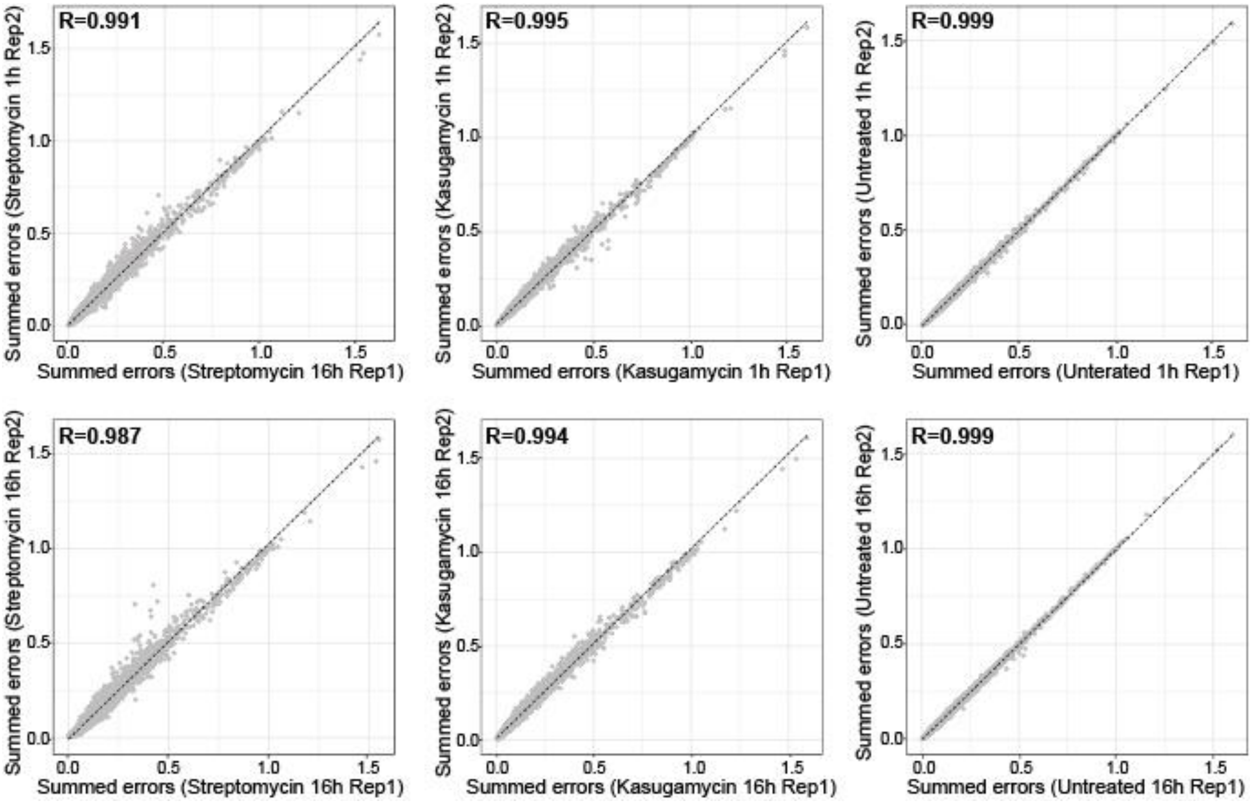

**Figure S20. Signal from all algorithms improves when using a WT matched dataset and data is normalized with a background comparison. (A)** Results obtained when comparing Streptomycin 1h Rep1 against WT 0h Rep1. **(B)** Results obtained when comparing Streptomycin 1h Rep1 against WT 1h Rep1.

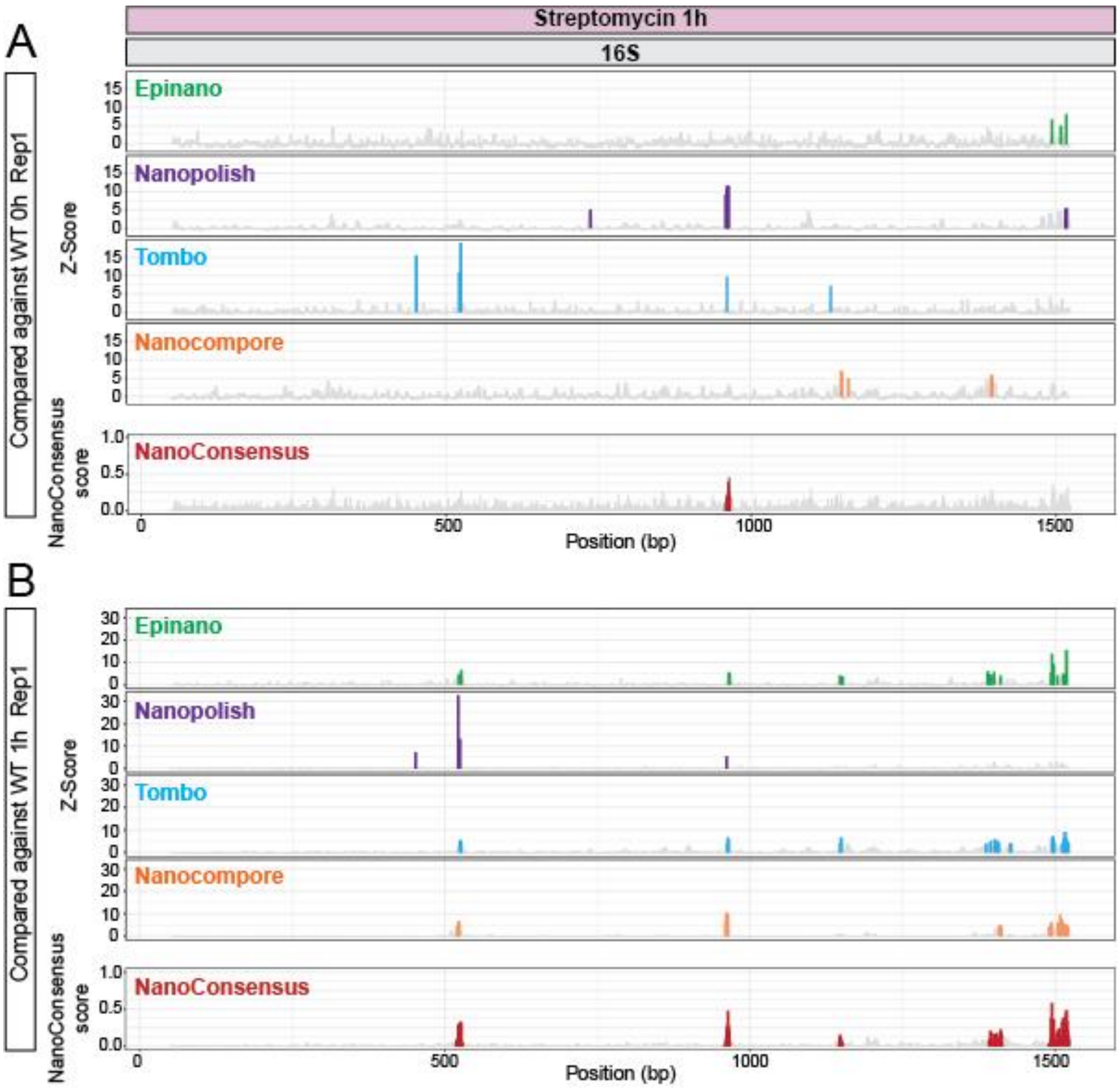

**Figure S21. Comparison of differential expression analysis using different types of alignments and counters. (A)** Differential expression analysis results between treated and untreated samples when aligning to the genome and using htseq-count with parameter *–nonunique all* to generate the per-read counts. In grey, non-significant genes; in green, genes with abs(log2FC) > 2 and; in red,  genes with abs(log2FC) > 2 and p adjusted value >=0.05. **(B)** Differential expression analysis results between treated and untreated samples when aligning to the transcriptome and using Salmon to generate per-transcript estimates. In grey, non-significant genes; in green, genes with abs(log2FC) > 2 and; in red,  genes with abs(log2FC) > 2 and p adjusted value >=0.05.

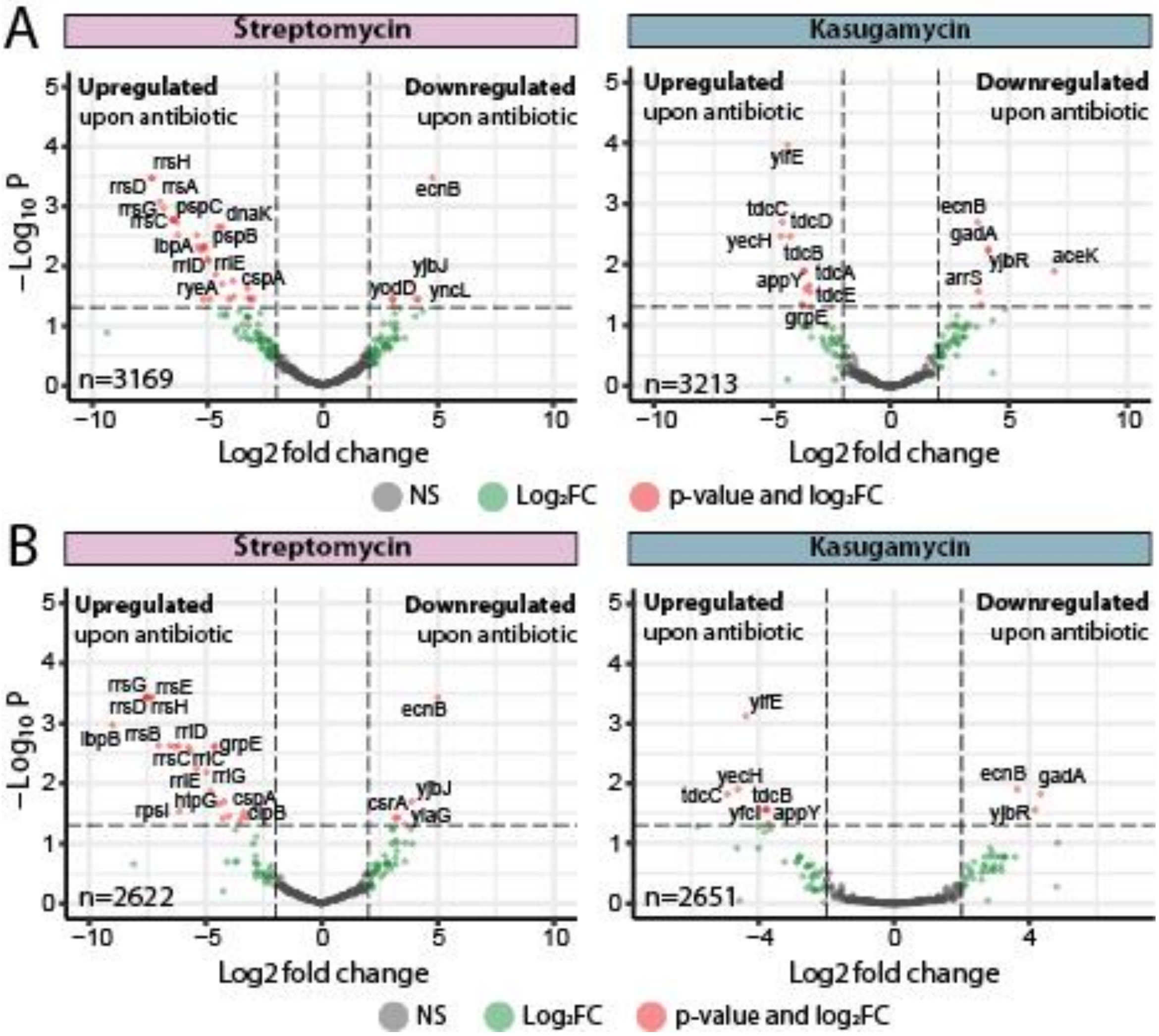
